# S18-phosphorylation of USP7 regulates interaction with TCEAL4 that defines specific complexes and potentially distinct functions

**DOI:** 10.1101/2021.07.07.451439

**Authors:** Francesca Querques, Sarah Darling, Izaak Cheetham-Wilkinson, Robbert Q Kim, Dharani K Hapangama, Titia K Sixma, Judy M Coulson

**Author notes:** Corresponding author: Judy M Coulson. Molecular Physiology and Cell Signalling, Institute of Systems Molecular & Integrative Biology, University of Liverpool, Liverpool, L69 3BX, UK. Cell Cycle Control Laboratory, UCL Cancer Institute, Paul O’Gorman Building, 72 Huntley St, London, WC1E 6HX, UK. Dept. of Cell and Chemical Biology, Leiden University Medical Centre, Leiden, The Netherlands (Robbert Q Kim).

## Abstract

USP7 is a nuclear deubiquitylase (DUB) with multiple cancer-associated substrates for which selective inhibitors are available, yet it remains unclear how the pleiotropic effects of USP7 are regulated. We report that S18-phosphorylation does not influence USP7 catalytic activity but instead confers selectivity for protein interactions. In particular, non-S18-phosphorylatable USP7 preferentially interacts with USP11 and TRIM27, together with TCEAL1 and TCEAL4 whose functions are unknown. Intriguingly, USP7 can interact with two cellular forms of TCEAL4, but USP11 only interacts with a lower abundance K142 mono-ubiquitylated form (TCEAL4-Ub), which can scaffold a complex containing both DUBs. Whilst USP11 and TCEAL4 are both USP7 substrates, TCEAL4-Ub levels are specifically maintained by USP11 with their levels positively correlated in cancer cell lines. Together these data illustrate how USP7 phosphorylation and TCEAL4 ubiquitylation combine to define distinct USP7 complexes. As TCEAL4 itself interacts with proteins involved in ubiquitylation and various forms of DNA regulation, these complexes may direct cellular activity of USP7.

## Introduction

Ubiquitylation is a complex post-translational modification (PTM) that regulates many cellular processes. Different configurations of mono-ubiquitin and poly-ubiquitin chains are appended to substrates by ubiquitin ligases, targeting proteins for degradation, or controlling signalling pathways through regulating protein activity or localisation ^1, 2^. Deubiquitylases (DUBs) can remove specific configurations of ubiquitin from defined substrates to reverse these processes ^3^ and have emerging functions within many cancer-associated pathways ^2, 4, 5^. Given their diverse substrates and complex roles, DUB activity must be tightly controlled within cells. A combination of PTMs, adaptor proteins and intramolecular rearrangement may regulate either DUB catalytic activity, or the recruitment of DUBs into specific complexes to direct their activity towards certain substrates ^6^.

Ubiquitin specific protease 7 (USP7, HAUSP) is a predominantly nuclear DUB with context-specific roles in cancer ^7^. It interacts with many other proteins ^8^ and has multiple established substrates, including the tumour-suppressors PTEN and p53 ^9, 10^ and oncogenes such as NOTCH1 and MDM2, a negative regulator of p53 ^11, 12^. The diverse USP7 substrates confer roles in the breadth of nuclear processes, from transcriptional and epigenetic regulation, through DNA replication, DNA repair and the cell cycle (reviewed in ^7, 13, 14^) as well as a cytoplasmic role in endosome recycling ^15^. These wide-ranging and sometimes opposing roles for USP7 necessitate tight control and precise direction of USP7 activity within cells. Structural studies show that USP7 exists in an inactive state, which can be switched to self-activate through intramolecular rearrangements that bring the C-terminal region into contact with the catalytic domain ^16, 17^; engagement with ubiquitylated substrate ^18^ and stabilisation of the active configuration by GMPS ^16, 19^ enhance catalytic activity.

USP7 has been described within several protein complexes that may modulate its catalytic activity or aid in recruitment to specific targets. Notably, many complexes partner USP7 with an E3 ligase, providing a switchable module that can either add or remove ubiquitin from a given substrate ^14^. In addition, tyrosine ^20^ or serine ^21^ phosphorylation may influence USP7 substrate preferences. Perhaps the best example of a USP7 substrate-switch is between deubiquitylating the E3 ligase MDM2 to promote p53 degradation, or deubiquitylating and thereby stabilising p53. DAXX promotes the interaction of USP7 with MDM2 ^22, 23^ whilst, in response to cellular stresses, cytoplasmic sequestration of GMPS is relieved allowing it to form a nuclear complex with USP7 that stabilises p53 ^24^. Reversible S18-phosphorylation of USP7 by CK2 and PPM1G has also been implicated in this switch during DNA damage responses, with dephosphorylation promoting p53 accumulation ^21^, although the mechanism remains to be elucidated.

Blocking USP7 activity can destabilise MDM2 to activate p53 and induce cell death ^11^ which, coupled with observations that USP7 overexpression is associated with poor prognosis in some cancers (reviewed in ^7^), highlight USP7 as an attractive drug target. Despite the limited specificity of first generation USP7 inhibitors, anti-tumorigenic activity was observed in some cancer models (reviewed in ^7^) and drug development programs have generated new lead compounds with high specificity for USP7 ^25, 26, 27, 28, 29^. However, as USP7 knockout mice exhibit embryonic lethality even in the absence of p53 ^30, 31^, alternative cellular roles of USP7 are clearly essential and these may underpin undesirable on-target effects of USP7 inhibition. Indeed, although recent studies suggest USP7 inhibitors predominantly suppress cancer cell growth through p53-dependent mechanisms ^29^, their applicability has been broadened to a sub-set of p53 mutant cancer cell lines and alternative predictive markers suggested ^28^. Thus, although USP7 is an attractive therapeutic target in cancer, we still need to better understand its context-specific regulation and substrates to rationally exploit USP7 inhibition.

To date, the only role described for S18-dephosphorylated USP7 is in DNA damage-dependent activation of p53 ^21^ suggesting it may act as a “stress response” DUB. However, the mechanism by which reversible S18-phosphorylation influences USP7 cellular behaviour, and hence its broader significance, remains unstudied. We investigated whether S18-status recruits USP7 to specific substrates or complexes and find that non-S18-phosphorylatable USP7 preferentially interacts with USP11, TRIM27, and two transcription elongation factor A (SII)-like (TCEAL) family members TCEAL1 and TCEAL4. We show that whilst both USP11 and TCEAL4 are USP7 substrates, USP7 can form a complex with TCEAL4 and TRIM27 independently of USP11. Intriguingly, USP11 only interacts with a minor cellular form of TCEAL4 that is K142 mono-ubiquitylated (TCEAL4-Ub). USP11 is required to maintain TCEAL4-Ub, and their levels are positively correlated in cancer cell lines. Importantly, TCEAL4-Ub scaffolds a complex containing both USP11 and USP7. We propose a model where S18-dephosphorylated USP7 and USP11 modulate the context and stability of TCEAL4, whilst TCEAL4 scaffolds a set of well-defined USP7 complexes. This interplay of PTMs and alternate complexes may help direct the cellular activity of USP7.

## Results

### S18-phosphorylation does not influence catalytic activity or gross cellular localisation of USP7

To assess how S18-phosphorylation may modulate USP7 behaviour, we examined whether it affected USP7 catalytic activity, either in biochemical assays, or within a cellular environment. Serine 18 was mutated in the context of full-length USP7, to create phospho-mimetic (S18E) or phospho-null (S18A) variants that were purified from bacteria together with wild-type (WT) USP7, which in this context is not phosphorylated (Supplementary Figure S1A). USP7 WT, S18A, or S18E all displayed *in vitro* activity towards ubiquitin-rhodamine (Figure 1A) with similar enzyme kinetics to those previously reported for full-length USP7 WT ^32, 33^, confirming this is unaffected by S18-status. To evaluate the cellular effects of S18-mutation, we generated stable U2OS cell lines that could be induced to express physiological levels of GFP-tagged WT, S18A, S18E, or catalytic-dead (C223S) USP7 (Supplementary Figure S1B). Within cells, WT and C223S GFP-USP7 exist as both S18-phosphorylated and dephosphorylated forms (Supplementary Figure 1C). We utilised the ubiquitin-VME (Ub-VME) suicide substrate to assess the on-rate of USP7 binding as an indication of DUB activity within cell lysates. Catalytic-dead C223S USP7, as expected, did not react with Ub-VME, however both S18A and S18E showed a similar reaction profile to WT GFP-USP7 (Figure 1B & 1C) consistent with the *in vitro* data. We also determined binding of endogenous USP7 to Ub-VME and found that this was comparable for S18-phosphorylated or non-S18-phosphorylated USP7 using phospho-specific antibodies (Figure 1D & 1E, Supplementary Figure S1D). USP7 is predominantly nuclear and, whilst catalytic-dead C223S GFP-USP7 partially re-distributed to the cytoplasm, S18A and S18E GFP-USP7 remained concentrated within the nuclear compartment in the majority of cells (Supplementary Figure S2). Together these data suggest that S18-status has no major impact on either generic DUB activity or gross cellular localisation of USP7, and likely affects USP7 behaviour in other ways.

**Figure 1.**
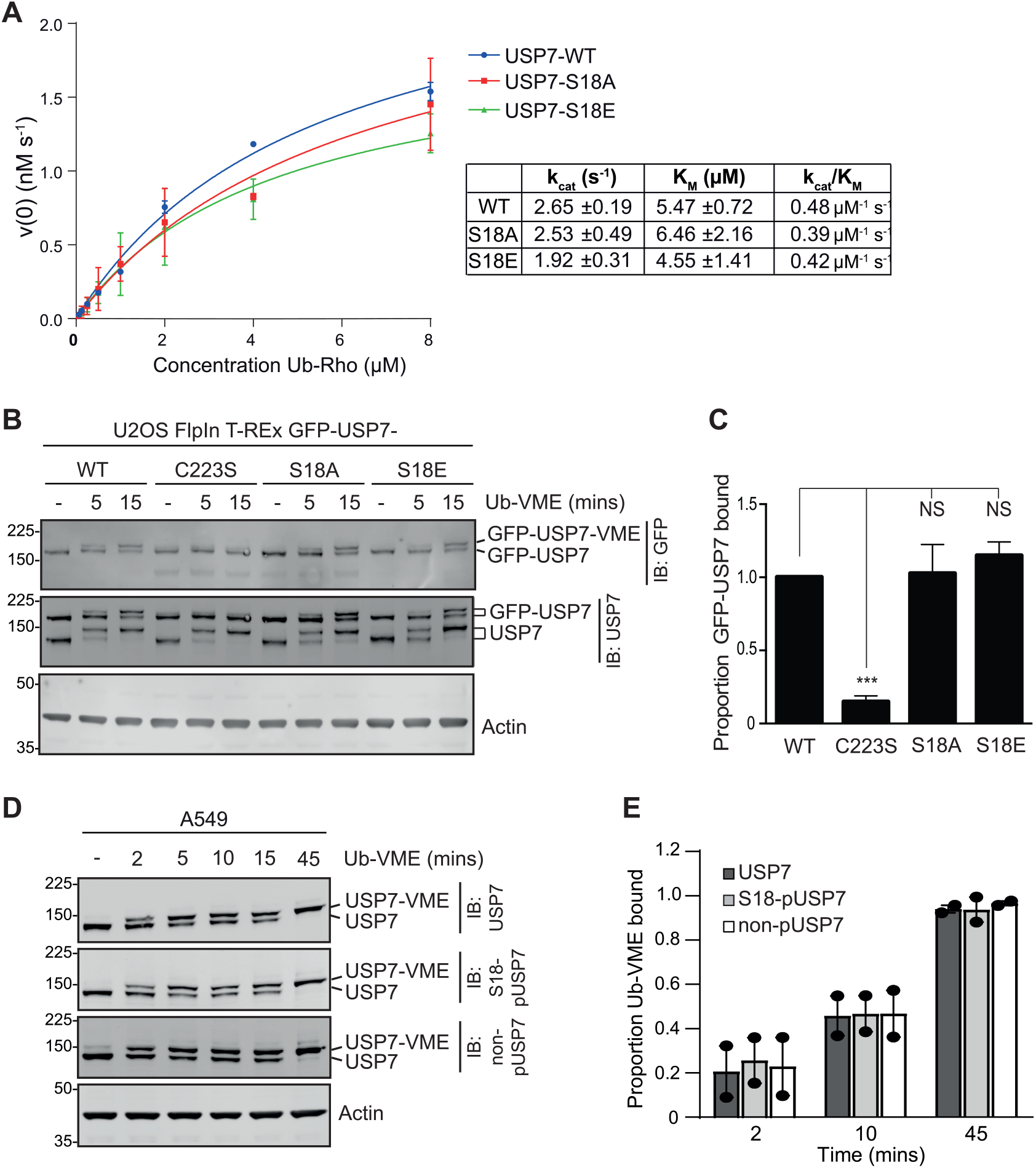
S18-status does not alter USP7 catalytic activity *in vitro* or *in vivo*. **A**, USP7 *in vitro* catalytic activity towards Ub-Rhodamine (Ub-Rho) is not affected by S18-mutation. Michaelis-Menten analysis of enzyme kinetics for recombinant full-length USP7-WT, S18A, or S18E proteins, experiments were conducted three times, using two independent protein preparations per construct. **B & C**, S18-mutation does not affect USP7 binding to Ub-VME. U2OS FlpIn T-REx GFP-USP7 cell lines were induced with doxycycline, equal amounts of lysates were incubated with Ub-VME and immunoblotted (**B**). Quantitative analysis of the proportion of GFP-USP7 bound to Ub-VME after 15 min (**C**); mean of 3 independent experiments, error bars SD, one-way ANOVA with Tukey’s post-hoc test; NS: not significant, ***p≤ 0.001. **D & E**, S18-phosphorylation does not affect USP7 binding to Ub-VME. A549 cell lysates incubated with Ub-VME and immunoblotted for S18-phosphorylated (S18-pUSP7), S18-unphosphorylated (non-pUSP7) or total USP7 to visualise endogenous USP7. Representative blot (**D**) and quantification of 2 independent experiments, error bars show range (**E**).

### S18-phosphomutants reveal selective protein interaction partners for USP7

Khoronenkova *et al*. previously demonstrated that S18-dephosphorylated USP7 had reduced ability to stabilise the E3 ligase MDM2, with concomitant accumulation of p53 ^21^. As our data (Figure 1) suggest this is not related to catalytic activity of USP7, we tested whether S18-status affects the interaction of USP7 with MDM2 or p53 by immunoprecipitation of GFP-USP7 variants from induced U2OS cells (Supplementary Figure S3). Catalytic-dead C223S USP7 demonstrated significant substrate-trapping of both MDM2 and p53. However, both S18 mutants pulled down similar amounts of MDM2 and p53 as WT USP7.

As USP7 has multiple substrates and interaction partners ^7^ we investigated whether S18-status affected the broader interaction profile of USP7. GFP-tagged USP7 WT, S18A or S18E were induced and immunoprecipitated from SILAC-labelled U2OS cells for mass spectrometry (Figure 2A, Supplementary Table 1) and 65 unique USP7 interactors were identified (Figure 2B, Supplementary Figure S4). Of these, 34 (52%) interacted with at least two USP7 variants, allowing quantification of relative pull-down dependent on S18-status. Unsupervised hierarchical clustering highlighted 10 of these proteins that show some selectivity for S18E USP7, and a smaller group of 4 proteins with S18A selectivity (Figure 2C). STRING analysis shows that these include several known USP7 interactors (Figure 2D). For example, the ubiquitin E3-ligase PJA1 and one of its substrates MAGED1 ^34^, which we show preferentially interact with phospho-mimetic USP7-S18E.

**Figure 2.**
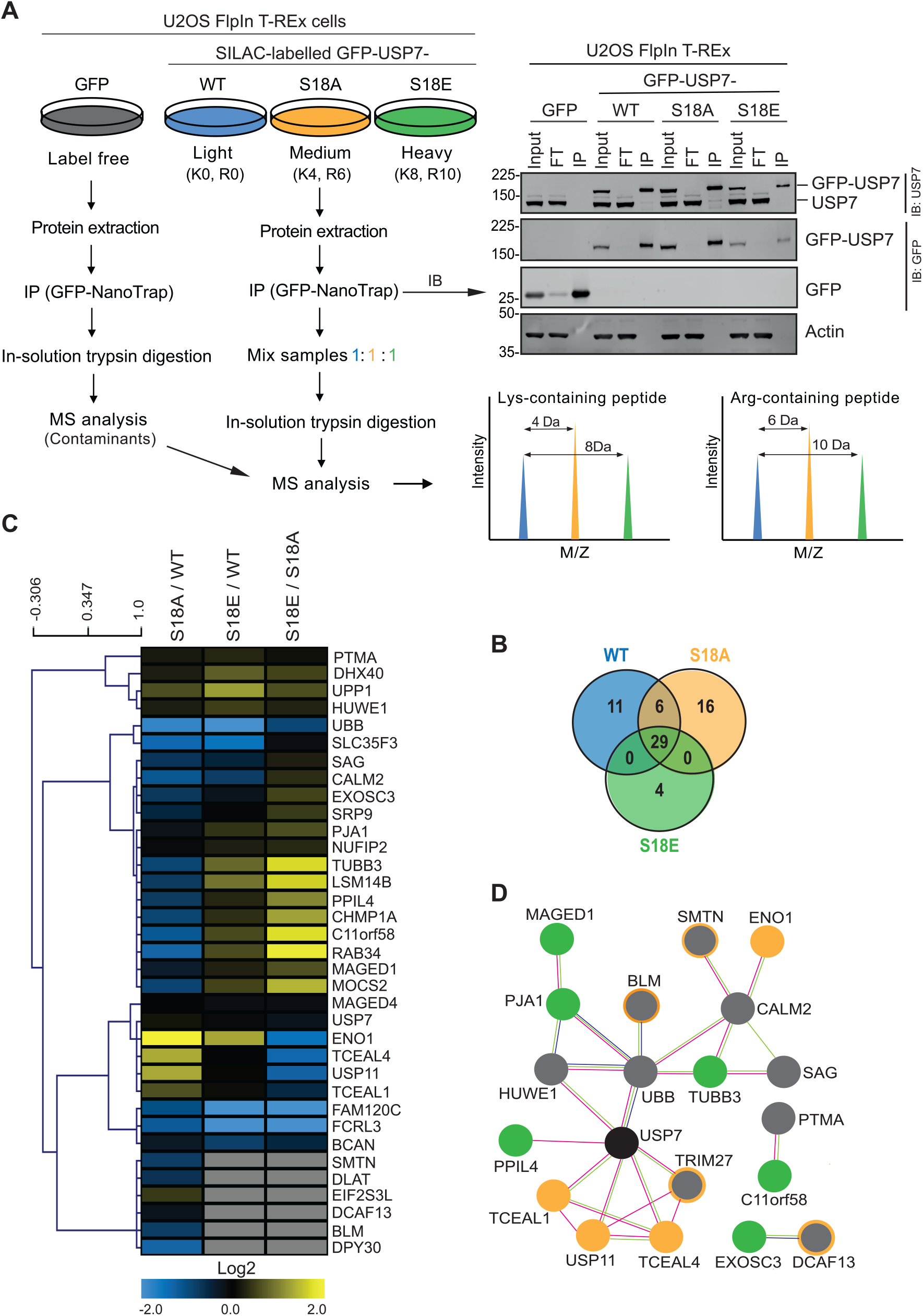
S18-phosphomutants reveal selective USP7 protein-protein interactions. **A**, Screen for differential protein interaction profiles dependent on USP7 S18-status. Schematic of the experimental strategy for SILAC labelling and GFP-USP7 pull-down in doxycycline-induced U2OS FlpIn T-REx GFP-USP7 cell lines. Lysates (1mg) were immunoprecipitated (IP) with GFP-NanoTrap beads. Unlabelled U2OS FlpIn T-REx GFP cells were used to filter out contaminants that bind GFP-beads. K: lysine, R: arginine, MS: mass spectrometry, FT: flow through. **B**, Venn diagram summarising proteins identified by mass spectrometry, excluding USP7 bait and contaminants. **C**, Unsupervised hierarchical clustering identifies proteins that differentially interact with USP7 S18-phosphomutants. Heatmap shows protein intensity ratios normalised to GFP-USP7 intensity in each sample. **D**, Network showing known and predicted protein-protein interactions from the STRING database. Interactors that preferentially bind S18A-USP7 (orange) or S18E-USP7 (green) are indicated (threshold: log2 S18E/S18A <-0.5 or >0.5). Grey circles, proteins with intensity below the threshold; grey circles with orange outline, interactors found only for S18A. Connections: known protein-protein interactions experimentally determined (pink) or from curated databases (blue), protein interactions or functional associations extracted from text mining (green).

Most notably, USP11 and TCEAL4, two high confidence interactors (HCIs) based on peptide coverage and intensity (Supplementary Figure S4), together with TCEAL1 and TRIM27, form a sub-network that selectively binds non-phosphorylatable S18A USP7 (Figure 2D). We verified this S18A-enriched network by directed co-immunoprecipitation, confirming that these four proteins preferentially interact with S18A compared to S18E or WT GFP-USP7 (Figure 3A & 3B, Supplementary Figure S5). Likewise, immunoprecipitation of endogenous USP11 selectively pulled down S18A USP7 (Figure 3C & 3D). Whilst TCEAL1 and TCEAL4 were previously reported as USP11 interactors ^8^, and TCEAL4 as a USP7 interactor ^35^, the significance of these interactions has not previously been addressed and our discovery that they preferentially form complexes with non-phosphorylatable USP7 is novel.

**Figure 3.**
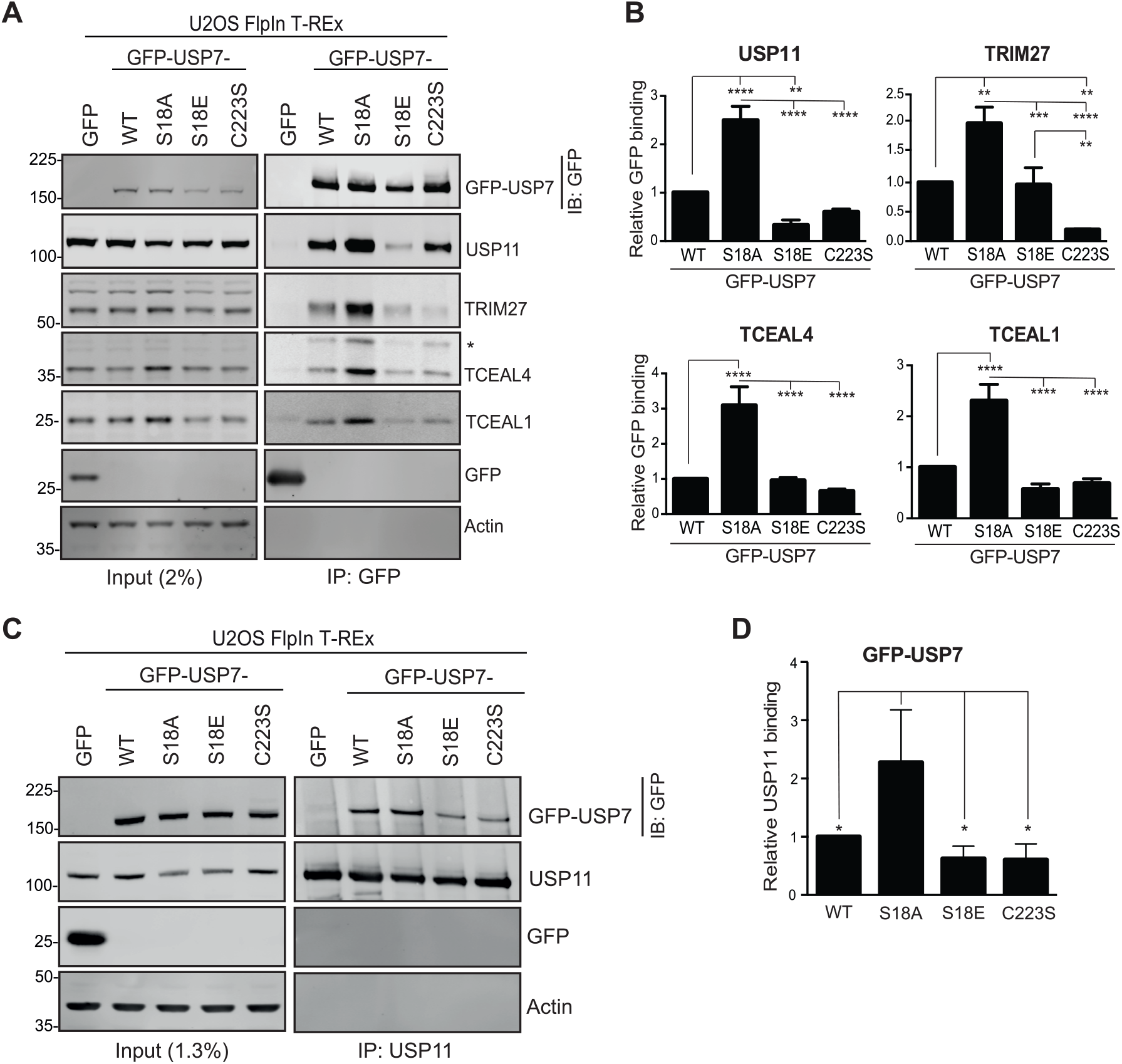
USP7-S18A preferentially interacts with USP11, TRIM27, TCEAL4 and TCEAL1. **A & B**, Directed immunoprecipitation confirms preferential S18A-USP7 interactors. U2OS FlpIn T-REx GFP-USP7 cell lines were induced with doxycycline and lysates subject to immunoprecipitation with GFP-NanoTrap beads. Representative immunoblot (**A**), *modified form of TCEAL4. Quantification of relative GFP-binding for interactors including the lower MW TCEAL4 species (**B**); mean values (normalised to GFP-USP7 in each IP) from 3 independent experiments, error bars SD. **C & D**, USP11 preferentially interacts with USP7-S18A. U2OS FlpIn T-REx were induced with doxycycline and lysates immunoprecipitated with USP11 antibody. Representative immunoblot (**C**) and quantification of relative USP11-binding for USP7 variants (**D**); mean values (normalised to USP11 in each IP) from 3 independent experiments, error bars SD. In **B & D**, one-way ANOVA with Tukey’s post-hoc test, *p≤ 0.05, **p≤ 0.01, ***p≤ 0.001, ****p≤ 0.0001.

### TCEAL4 associates with other complexes that may infer cellular roles

The TCEAL family are named by similarity to TFIIS/TCEA, which is involved in transcription elongation and transcript fidelity, although little is currently known about their functions. They are encoded within an expansion of rapidly evolving genes on human chromosome Xq22.1 that includes the *BEX, GASP* and *TCEAL* (*WEX*) gene families ^36^. Phylogenetic analysis of the human TCEAL family proteins (Supplementary Figure S6A) shows two major clusters that segregate TCEAL4 and TCEAL1. TCEAL4 was the higher confidence USP7 interactor (Supplementary Figure S4) and shares 78% identity with its closest paralog TCEAL2 (Supplementary Figure S6B). The *TCEAL* genes are absent in lower eukaryotes and, whilst *TCEAL2* and *TCEAL4* are also not present in mice ^36, 37^, TCEAL4 is well-conserved across primates (Supplementary Figure S6C). The TFA (BEX) domain is conserved amongst the TCEAL and BEX protein families, although its function is unclear. TCEAL4 has a predicted bipartite nuclear localisation signal and multiple phosphorylation sites, most notably for CK2, whilst ubiquitylation and SUMOylation of TCEAL4 are reported in high throughput studies collated at Phosphosite ^38^ (Supplementary Figure S6D).

Mechanistic roles are not well-defined for the majority of the TCEAL family, perhaps best characterised to date is TCEAL7, which can influence MYC and NFkB-driven transcription ^39, 40^. We confirmed that GFP-TCEAL4 predominantly localises in the nucleus of U2OS cells (Supplementary Figure S6E) and thus may potentially be involved in nuclear processes such as transcription. However, no TCEAL4 functions have yet been described, and the related BEX proteins are suggested to be hubs that participate in multiple cellular pathways ^37^. To investigate further, we immunoprecipitated GFP-TCEAL4 for mass spectrometry (Figure 4A, Supplementary Table 2) and identified several hundred interacting proteins, substantially increasing the list of TCEAL4 interactors collated from high-throughput studies (Figure 4B). USP7 and USP11 were two of the strongest HCIs based on peptide coverage and intensity, whilst TRIM27 was detected with lower confidence (Figure 4C). Directed immunoprecipitation experiments of GFP-TCEAL4 further verified its interaction with endogenous USP7, USP11 and TRIM27, as well as TCEAL1, even though the latter was not identified by mass spectrometry (Figure 4D). The broader TCEAL4 interactome was enriched for proteins involved in cellular responses to external stimuli and stress, tRNA-aminoacylation, the cell cycle and multiple components of the ubiquitin proteasome system (UPS) (Figure 4E & 4F); proteins involved in DNA-related processes including DNA replication and repair were also well represented (Figure 4F & 4G). Together these data suggest that TCEAL4 may play diverse cellular roles. Several TCEAL4 HCIs have previously been identified as interactors of USP7 or USP11, and we speculate that the processes to which these interactors map may suggest alternative functions for USP7 in complex with TCEAL4 and/or USP11 (Figure 4H). Notably, the 18 common interactors of TCEAL4, USP7 and USP11 form an interconnected network that includes USP15, E3 ligases and proteins associated with mitosis (Figure 4I), potentially inferring specific cellular functions for the TCEAL4/USP7/USP11 complex.

**Figure 4.**
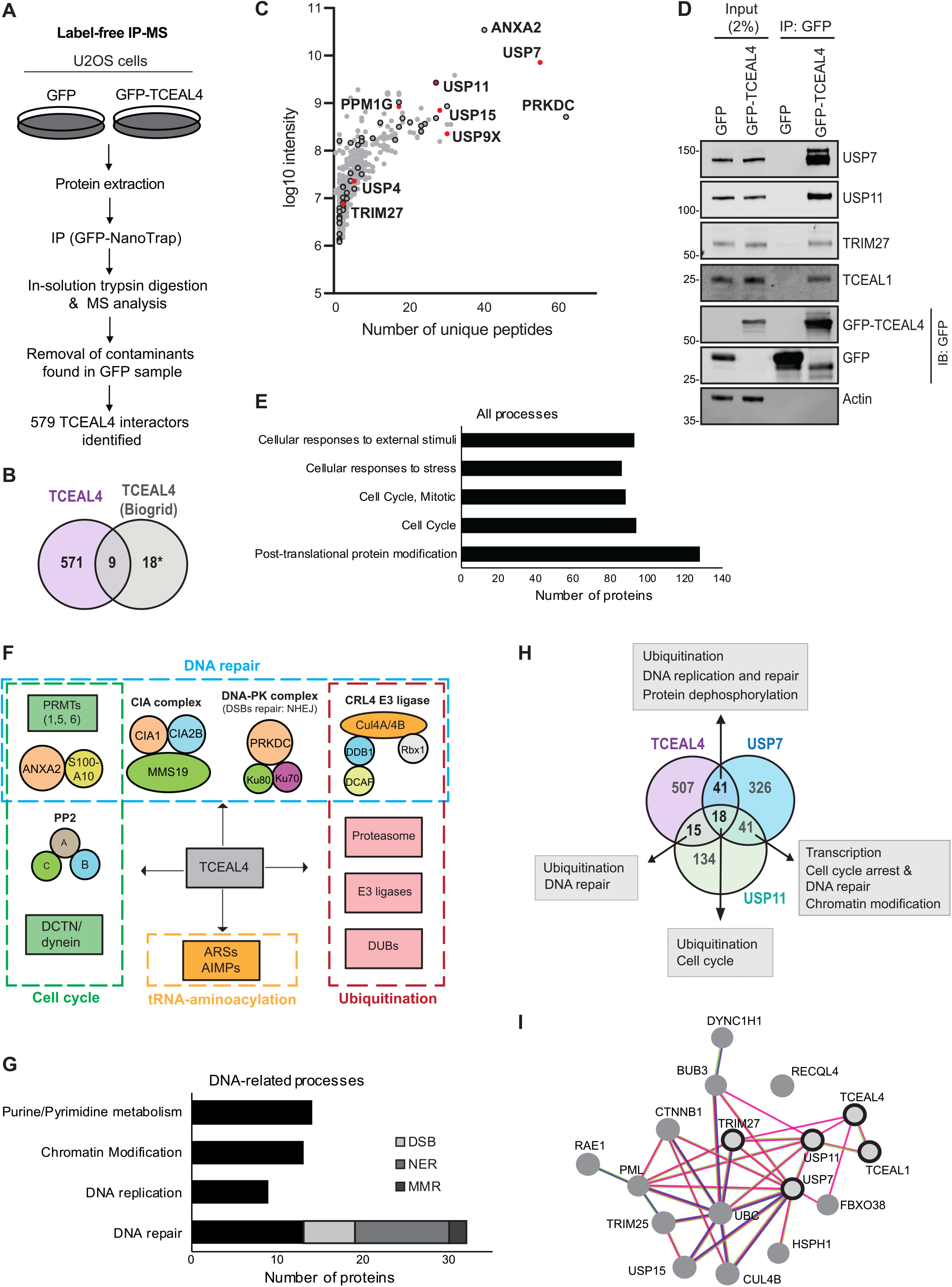
TCEAL4 protein interactions infer cellular roles. **A**, Schematic of label-free affinity mass spectrometry strategy to identify TCEAL4 interacting proteins. U2OS cells were transfected for 24 hrs with a plasmid encoding GFP alone or GFP-tagged TCEAL4 WT isoform-1, and lysates were immunoprecipitated with GFP-NanoTrap beads. **B**, Venn diagram shows the intersection of TCEAL4 interactors identified in this study by MS plus TCEAL1 (confirmed by directed IP), with the interactors reported in Biogrid for TCEAL4. *3 interactors reported in Biogrid were also present in our dataset but did not satisfy the cut-off criteria (TCEAL2, no unique peptides identified; NEDD1 and USP5, detected in GFP sample with intensity ratio TCEAL4/GFP <4). In all other panels, TCEAL4 interactors are those identified in this study by MS and/or directed IP. **C**, Scatter plot summarising identified TCEAL4 interactors after the removal of contaminants present in the GFP sample. Red indicates DUBs and other proteins of interest; black borders indicate proteins involved in DNA repair. **D**, GFP-TCEAL4 interacts with USP7, USP11, TRIM27 and TCEAL1. U2OS cells were transfected for 24 hrs with a plasmid encoding GFP or GFP-TCEAL4; lysates were immunoprecipitated with GFP-NanoTrap beads and representative immunoblots from 2 independent experiments are shown. **E**, TCEAL4 interactors were interrogated using Gene Set Enrichment Analysis (GSEA) with the molecular signatures database (MSigDB) to identify enriched pathways; histogram represents the top 5 most significantly enriched pathways. **F**, Schematic illustrating protein complexes or functionally-related protein groups identified through manual annotation, where at least one protein was in the top 20% of TCEAL4 interactors based on intensity. **G**, TCEAL4 interactome for DNA-related processes. TCEAL4 interactors were interrogated for proteins in indicated Reactome/KEGG pathways (GSEA/MSigDB). Proteins unequivocally belonging to a single DNA repair process were assigned: DSB, double-strand break repair (non-homologous end-joining (NHEJ) and homologous recombination (HR)); NER, nucleotide excision repair; MMR, mismatch repair. **H**, TCEAL4 may scaffold USP7/USP11 complexes with distinct functions. Venn diagram shows the intersection of TCEAL4 interactors with interactors reported in Biogrid for USP7 and USP11. **I**, String analysis highlights the interconnected proteins that can interact with TCEAL4, USP7 and USP11 (centre of Venn in **G**) extending upon our initial network (black outlines); FBXO38 interaction added from ref ^48^. Connections: known protein-protein interactions experimentally determined (pink) or from curated databases (blue), protein interactions or functional associations extracted from text mining (green).

### Cellular TCEAL4 exists as two different species

Intriguingly we noticed that two TCEAL4 immunoreactive species are evident in U2OS cells, both of which preferentially interact with S18A USP7 (Figure 3A & Supplementary Figure S5) and are depleted by TCEAL4 siRNAs (Figure 5A). The major form migrates at around 36kDa, whereas a less abundant 42kDa form is stabilised by proteasome inhibition with epoxomicin (Figure 5A & 5B). We excluded the possibility that these might result from cross-detection of the TCEAL2 paralog, as no unique TCEAL2 peptides were identified in the USP7 interactome, whilst two TCEAL4 antibodies and three of the four TCEAL4 siRNAs do not cross-react with TCEAL2 (Supplementary Figure S7). We also investigated whether they could represent alternative TCEAL4 isoforms. The predicted molecular weights for TCEAL4 isoform-1 (NP_079139, Q96EI5-1, 215aa) and isoform-2 (NP_001287830, Q96EI5-2, 358aa) are 25kDa and 40kDa, respectively. As no 25kDa form is evident in U2OS cells (Supplementary Figure S8A) we wondered whether they might predominantly express isoform-2, however only TCEAL4 isoform-1 transcripts were detected by qRT-PCR (Supplementary Figure S8B). Furthermore, exogenous MYC/FLAG-tagged TCEAL4 isoform-1 also migrated as two species, whose abundance and apparent molecular weight correspond to those of endogenous TCEAL4 isoform-1 (Supplementary Figure S8C).

**Figure 5.**
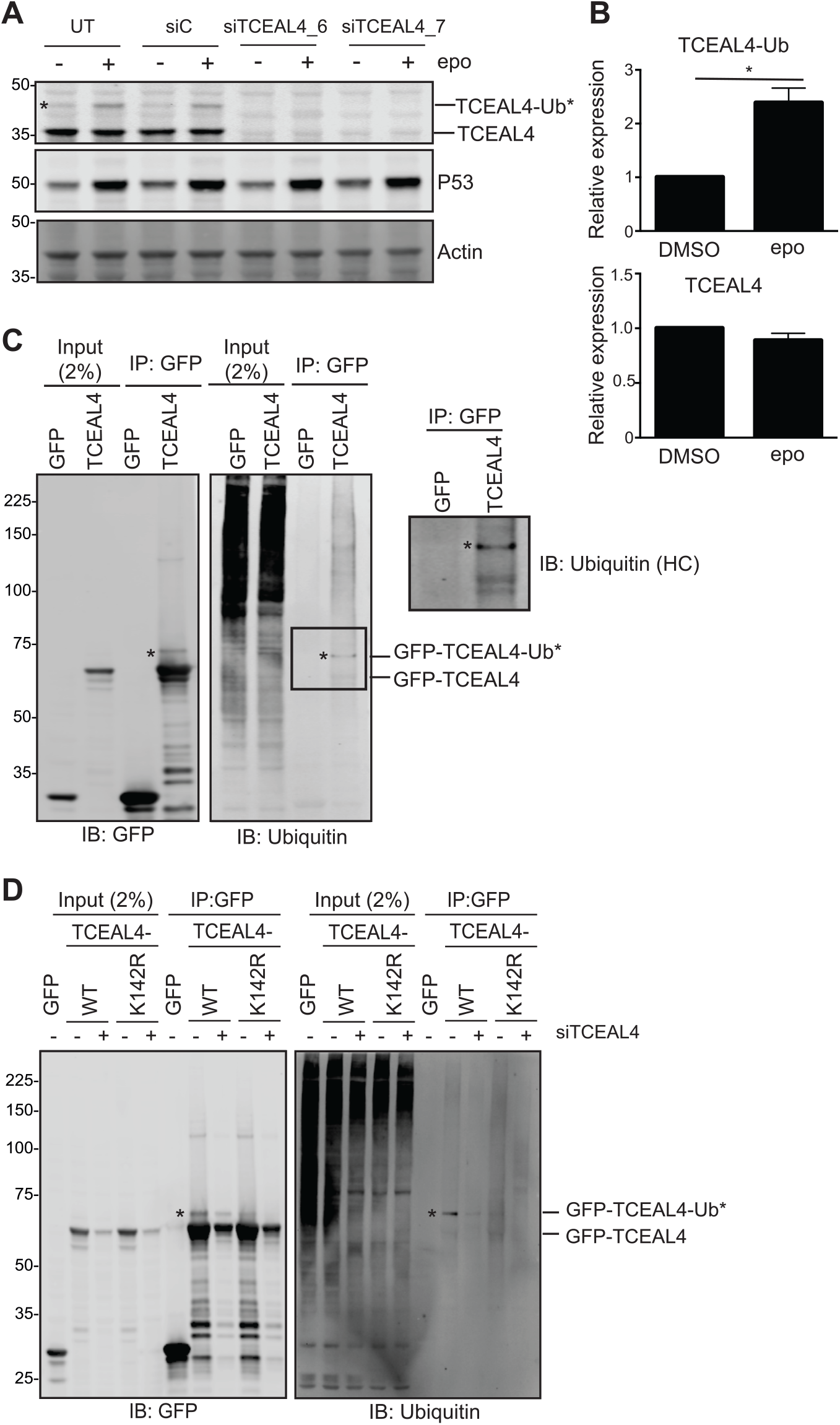
TCEAL4 is subject to post-translation modification that affects stability. **A & B**, A higher MW TCEAL4-specific immunoreactive band increases upon proteasomal inhibition. Uninduced U2OS FlpIn T-REx S18A-USP7 cells, either untransfected (UT) or transfected for 72 hrs with control (siC) or TCEAL4 siRNAs, were treated with DMSO or epoxomicin (epo) for 6 hrs; representative immunoblot includes p53 as a control for proteasome inhibition (**A**). Quantification of untransfected samples (**B**); mean values (normalised to actin and expressed relative to siC) from 3 independent experiments, error bars SD, one-sample Student’s t-test, *p≤ 0.05. **C**, TCEAL4 can be mono-ubiquitylated. U2OS cells were transfected for 24 hrs with a plasmid encoding GFP or GFP-TCEAL4 and treated with epoxomicin for 6 hrs; representative immunoblot from 3 independent experiments for lysates immunoprecipitated with GFP-NanoTrap beads. **D**, K142 is a major site of TCEAL4 mono-ubiquitylation. U2OS cells, either untransfected or transfected with siTCEAL4_7 for 72 hrs, were transfected with plasmids encoding GFP or GFP-tagged TCEAL4 WT or K142R for the final 24 hrs and treated with epoxomicin for the final 6 hrs; representative immunoblot from 3 independent experiments for lysates immunoprecipitated with GFP-NanoTrap beads.

Together these data suggest there are two species of TCEAL4 isoform-1, and we explored whether these might reflect post-translational modifications. As phosphatase treatment did not alter migration of either TCEAL4 species (Supplementary Figure S8D) we speculated that TCEAL4 may be modified by ubiquitylation or SUMOylation. We immunoprecipitated GFP-TCEAL4 isoform-1 from transiently transfected U2OS cells and probed for ubiquitin or SUMO-1. The more abundant lower molecular weight (MW) species (consistent with endogenous 36kDa TCEAL4) was immunoreactive for SUMO-1 (Supplementary Figure S8E) but, although global studies have identified K29 or K142 SUMOylation ^41, 42, 43^, we were unable to confirm this by mutagenesis of K29, K142, or six other predicted SUMO consensus sites (Supplementary Figure S6D, Supplementary Figure S8F & S8G). In contrast, the less abundant higher MW species (consistent with endogenous 42kDa TCEAL4) was not SUMO-1 immunoreactive but was instead recognised by a ubiquitin antibody (Figure 5C). Global ubiquitylation studies have identified modification of TCEAL4 at K139, K142 and K164, with K142 most commonly reported (https://www.phosphosite.org; Supplementary Figure S6D). Indeed, K142R mutation led to selective loss of the higher MW GFP-TCEAL4 species (Figure 5D, Supplementary Figure S8F) confirming mono-ubiquitylation at this residue. Henceforth, this modified species is referred to as TCEAL4-Ub, to distinguish it from the more abundant lower MW protein referred to as TCEAL4.

### The two TCEAL4 species define distinct cellular USP7 or USP11 complexes

Immunoprecipitation of endogenous USP11 confirmed its cellular interaction with USP7, TCEAL1 and TCEAL4-Ub, however we found that USP11 cannot interact with TCEAL4 (Figure 6A). We also expressed MYC/FLAG-tagged TCEAL4 isoform-1 in U2OS cells to verify this observation, and again only the TCEAL4-Ub species co-immunoprecipitated with USP11 (Figure 6B). Furthermore, although GFP-TCEAL4 pulled-down all components of the endogenous complex, GFP-TCEAL4 bearing the K142R mutation, which ablates the primary ubiquitylation site, impeded pull down of USP11 (Supplementary Figure S9A). Together these data confirm that USP11 is selectively incorporated into complexes that contain TCEAL4-Ub and, remarkably, that USP11 does not appear to remove the mono-ubiquitin from TCEAL4-Ub.

**Figure 6.**
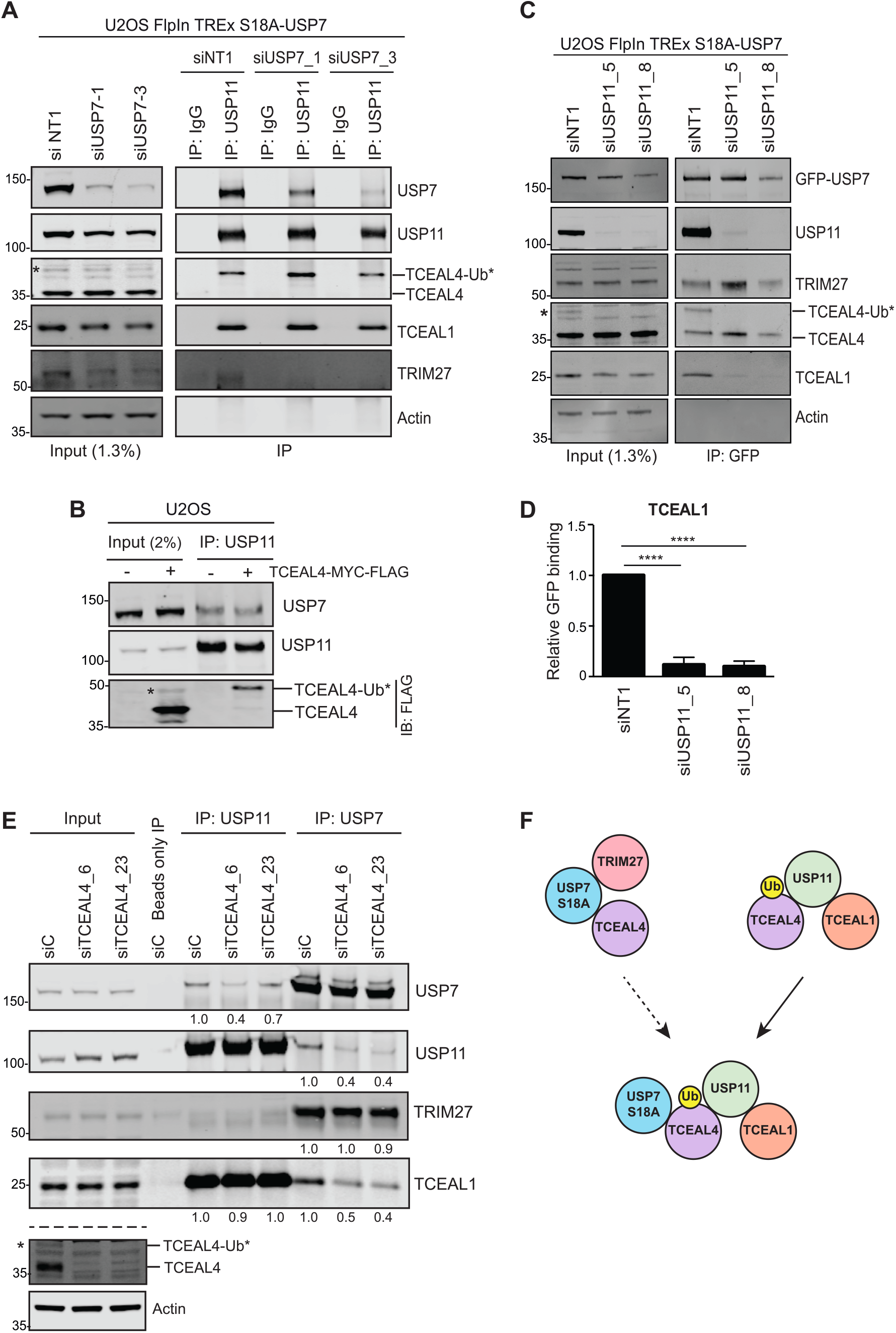
TCEAL4 species define different S18A-USP7-enriched protein complexes. **A**, USP7 is required for USP11 interaction with TRIM27, but not with TCEAL4-Ub or TCEAL1. Uninduced U2OS FlpIn T-REx S18A-USP7 cells were transfected with control (siNT1) or USP7 siRNAs for 72 hours, and lysates were immunoprecipitated with an anti-USP11 antibody; representative immunoblot from 3 independent experiments. **B**, USP11 specifically interacts with TCEAL4-Ub. U2OS cells were transfected for 24 hrs with pCMV6-TCEAL4-MYC-FLAG and treated with epoxomicin for 6 hrs. Lysates were immunoprecipitated with an anti-USP11 antibody; representative immunoblot from 2 independent experiments. **C**, USP11 is required for USP7 interaction with TCEAL4-Ub and TCEAL1. U2OS FlpIn T-REx S18A-USP7 cells were transfected with control (siNT1) or USP11 siRNAs for 72 hours and induced with doxycycline for the last 24 hours. Lysates were immunoprecipitated with GFP-NanoTrap beads; representative immunoblot from 3 independent experiments. **D**, USP11 depletion abolishes TCEAL1 interaction with USP7. Quantification of IP from (**C**); mean values from 3 independent experiments (normalised to the amount of GFP-USP7 pulled-down in each sample); error bars SD, one-way ANOVA with Tukey’s post-hoc test, ****p≤ 0.0001. **E**, TCEAL4 depletion decreases interaction of USP7 with USP11. U2OS cells were transfected with control (siC) or TCEAL4-specific siRNAs for 72 hrs and lysates subject to immunoprecipitation with USP7 or USP11 antibodies. Input: 1% USP7 IP, 1.4% USP11 IP. Representative immunoblot, numbers indicate mean expression from 2 independent experiments (normalised to USP7 or USP11 pull-down and expressed relative to siC). **F**, TCEAL4 species participate in alternative USP7 or USP11 complexes.

To further explore whether assembly of TCEAL4 complexes is dependent on specific DUBs, we tested the effect of USP7 or USP11 depletion on protein interactions in U2OS cells. First, we immunoprecipitated endogenous USP11 from USP7-depleted samples. The interaction of USP11 with TCEAL1 or TCEAL4-Ub was not affected, however USP11 did not interact with TRIM27 in the absence of USP7 (Figure 6A). Consistent with that observation, GFP-TCEAL4 did not pull down TRIM27 in USP7-depleted cells (Supplementary Figure S9B). Thus, USP11 can exist in an independent complex with TCEAL1 and TCEAL4-Ub, whilst USP7 is required to recruit TRIM27 (Figure 6F). We then depleted USP11 from U2OS cells and found that TCEAL4-Ub was not detected in the input samples (Figure 6C) indicating that USP11 is necessary for its presence. When endogenous USP7 was immunoprecipitated from these USP11-depleted samples, TRIM27 and TCEAL4 but not TCEAL1 were pulled down (Figure 6C, Figure 6D). This suggests that USP7 does not directly interact with TCEAL1 and can exist in an independent complex with non-ubiquitylated TCEAL4 and TRIM27 (Figure 6F).

### TCEAL4-Ub scaffolds USP7 and USP11 interaction

We speculated that the different TCEAL4 species may be important in formation of distinct USP7 complexes. Indeed, depletion of TCEAL4 in U2OS cells substantially reduced the interaction of USP7 with both USP11 and TCEAL1 (Figure 6E) suggesting it promotes interaction of the two DUBs. Intriguingly, no endogenous TCEAL4-Ub is present in HCT116 cells and here GFP-USP7 pulled down TCEAL4 but little USP11 (Supplementary Figure S10A), whilst no interaction between endogenous USP7 and USP11 was evident (Supplementary Figure S10B) suggesting the USP7/TCEAL4 complex is predominant (Supplementary Figure S10C). Importantly, GFP-TCEAL4 overexpression restored robust interaction of endogenous USP11 and USP7 in HCT116 cells (Supplementary Figure S10D). This appears to be mediated by TCEAL4-Ub, as GFP-TCEAL4 can be ubiquitin-modified in HCT116 and pulled down both endogenous USP7 and USP11, whilst the GFP-TCEAL4-K142R ubiquitylation mutant pulled down less USP11 (Supplementary Figure S10E). Taken together these data support the idea that TCEAL4-Ub can scaffold a larger complex containing both DUBs (Figure 6F).

### USP11 and TRIM27 are stabilised by USP7 in U2OS cells

Our next question was whether any components of these complexes were substrates of either DUB. Indeed, experiments in U2OS indicated that components of the S18A-selective interaction complex may be stabilised by USP7. TRIM27 levels were markedly decreased by USP7 depletion (Figure 6A left panel, Supplementary Figure S11A) consistent with a previous report that USP7 protects TRIM27 from auto-ubiquitylation and proteasomal degradation ^15^. USP7 depletion also significantly reduced USP11 levels (Figure 6A left panel, Supplementary Figure S11A) and increased USP11 ubiquitylation (Supplementary Figure S11B) suggesting that USP7 stabilises USP11 by deubiquitylation in U2OS cells. Interestingly however, USP7 depletion did not destabilise USP11 in HCT116 cells (Supplementary Figure S10F) that lack TCEAL4-Ub and where interaction of these endogenous DUBs was not evident. These observations are consistent with a previous study that proposed a context-dependent role for USP7 in regulating USP11 ^35^. Our data suggest that TCEAL4-Ub mediates an interaction between USP7 and USP11, which allows USP7 to act upon USP11 as substrate and protect it from degradation.

### USP7 stabilises TCEAL4 whilst USP11 stabilises TCEAL4-Ub

Depletion of either USP7 or USP11 in U2OS cells reduced the levels of TCEAL1 and of specific TCEAL4 species (Figure 6A & 6C, left panels; Supplementary Figure S11A & S11C). The level of the more abundant lower MW TCEAL4 species was reduced by USP7 depletion, although TCEAL4 transcript expression was not affected (Supplementary Figure S11D). In HCT116 cells, where USP7 and TCEAL4 also interact (Supplementary Figure S10A & S10E), TCEAL4 was modestly reduced by USP7 depletion (Supplementary Figure S10F). In contrast, TCEAL4-Ub was only detected in U2OS cells in the presence of USP11 with which it specifically interacts (Figure 6C left panel & Supplementary Figure S11C). Following USP11-depletion we found that TCEAL4 accumulated, suggesting that TCEAL4-Ub is converted to TCEAL4 in the absence of USP11 (Supplementary Figure S11E). Supporting this, TCEAL4 did not accumulate in USP11-depleted HCT116 cells that lack endogenous TCEAL4-Ub (Supplementary Figure S10F).

TCEAL4-Ub appears to be unstable in U2OS cells as proteasome inhibition led to its accumulation (Figure 5A & 5B), but TCEAL4-Ub levels were not rescued by epoxomicin in the absence of USP11 (Figure 7A) presumably because of its conversion to TCEAL4. This conversion relies on TCEAL4 stabilisation by USP7, as co-depleting USP7 and USP11 induced poly-ubiquitylation of GFP-TCEAL4 (Figure 7B) and proteasome inhibition was now required to accumulate TCEAL4 (Figure 7A). As expected, inhibition of USP7 DUB activity with XL177A also reduced TCEAL4 but had little effect on TCEAL4-Ub (Figure 7C, Supplementary Figure S12). Furthermore, as observed with USP7 siRNAs, epoxomicin was required to accumulate TCEAL4 from TCEAL4-Ub when USP7 inhibition with XL177A was combined with USP11 depletion (Figure 7C). Our working model (Figure 7D) proposes that TCEAL4-Ub is either generated or protected within a USP11 complex, where USP11 cannot remove the mono-ubiquitin. This can drive the formation of a larger complex with USP7, which may be reinforced through stabilisation of USP11 by USP7, and where TCEAL4-Ub remains protected. However, in the absence of USP11, TCEAL4-Ub is rescued from degradation through deubiquitylation by USP7, which under these conditions appears to also remove the mono-ubiquitin. Therefore, only in the absence of both USP7 and USP11 activity does TCEAL4-Ub become poly-ubiquitylated and degraded.

**Figure 7.**
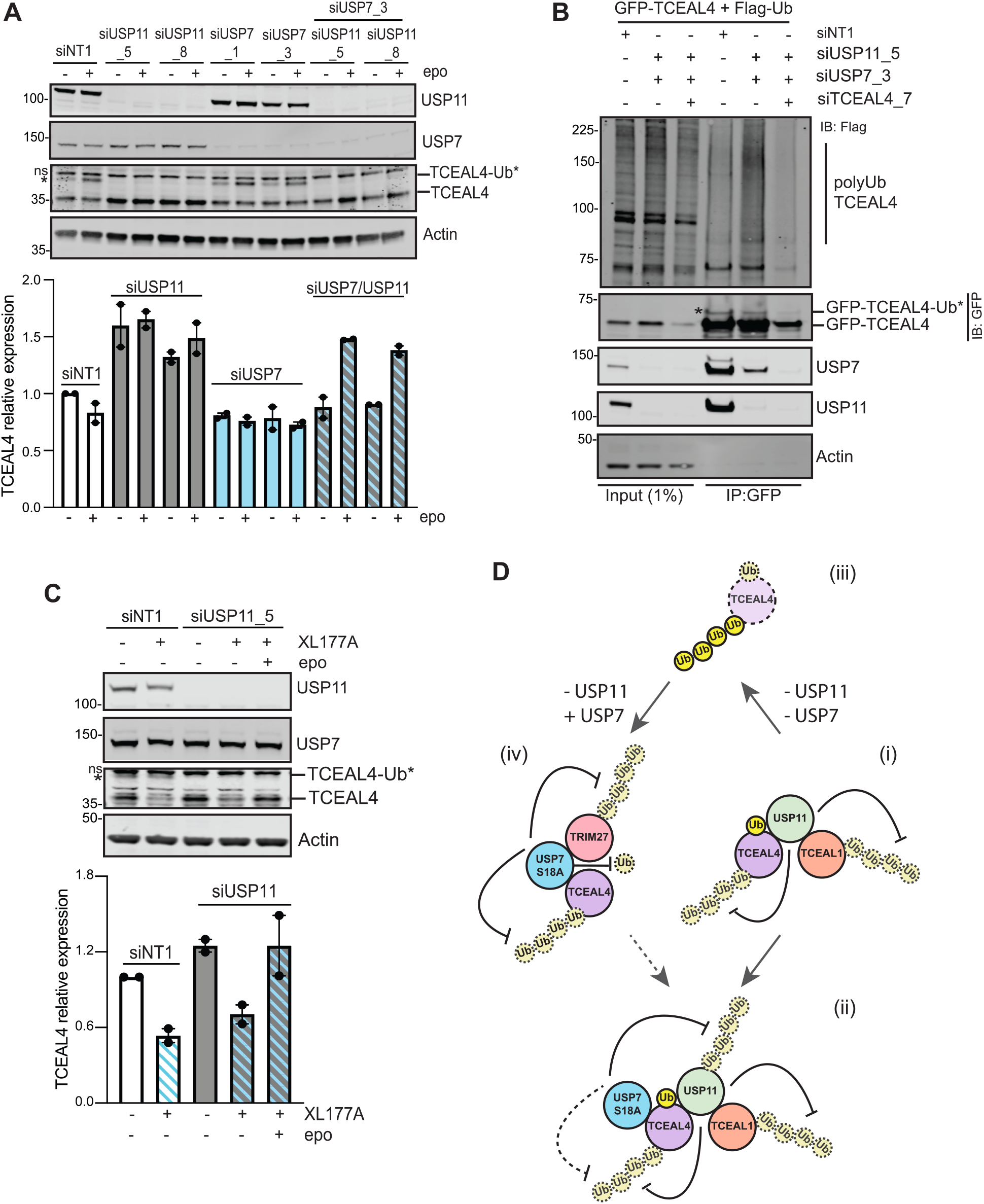
USP7 is required to accumulate TCEAL4 and USP11 is required to maintain TCEAL4-Ub. U2OS cells were transfected with siRNAs for 72 hours, co-transfected with plasmids for the final 24hrs, and treated with DMSO, epoxomicin (epo) and/or XL177A for 6 hrs, as indicated. **A**, USP7 is required to accumulate TCEAL4 in the absence of USP11. A representative immunoblot and mean expression of TCEAL4 from 2 independent experiments, normalised to actin and relative to siNT1 in DMSO, error bars show range. ns, non-specific. **B**, TCEAL4 becomes poly-ubiquitylated in the absence of USP7 and USP11. Lysates were immunoprecipitated with GFP-NanoTrap beads; representative immunoblot from 2 independent experiments. **C**, Inhibition of USP7 deubiquitylase activity blocks TCEAL4 accumulation. A representative immunoblot and mean expression of TCEAL4 from 2 independent experiments, normalised to actin and relative to siNT1 in DMSO; error bars represent range. **D**, Proposed model by which USP7 and USP11 stabilise alternative forms of TCEAL4. The less stable TCEAL4-Ub form is protected or generated within the USP11 complex (i) and in the larger complex (ii) this may be reinforced by USP7 either directly or via deubiquitylation of USP11. However, in the absence of USP11, TCEAL4-Ub can be poly-ubiquitylated and degraded (iii) unless it is rescued by USP7 deubiquitylation and stabilised as the TCEAL4 form (iv). Poly-ubiquitylation of TCEAL4 is indicated independently to K142 mono-ubiquitylation for simplicity, although this may also include ubiquitin chain extension at K142.

### TCEAL4 is a candidate tumour suppressor and its USP7 complexes may be dysregulated in cancer

In normal tissues, TCEAL4 gene expression is highest in female reproductive organs, especially in the ovaries and uterus (Supplementary Figure S13A). TCEAL4 is frequently downregulated in ovarian and endometrial cancers, suggesting it acts as a tumour suppressor (Figure 8A, Supplementary Figure S13B-S13D). In the CPTAC endometrial cancer proteogenomics dataset ^44^ TCEAL4 protein expression was significantly reduced in tumours relative to age-matched normal controls (Figure 8B). There was also a trend for USP11 protein downregulation in tumours within this study (Figure 8C) whilst USP7 protein expression, and more markedly S18-phosphorylated USP7, were higher in tumour tissue (Figure 8D & 8E). These data suggest that TCEAL4/S18-dephosporylated USP7 complexes could be modulated in cancer through multiple mechanisms, including downregulation of TCEAL4 gene expression, as well as via reduced USP11 levels or upregulated USP7 phosphorylation (Figure 8F).

**Figure 8.**
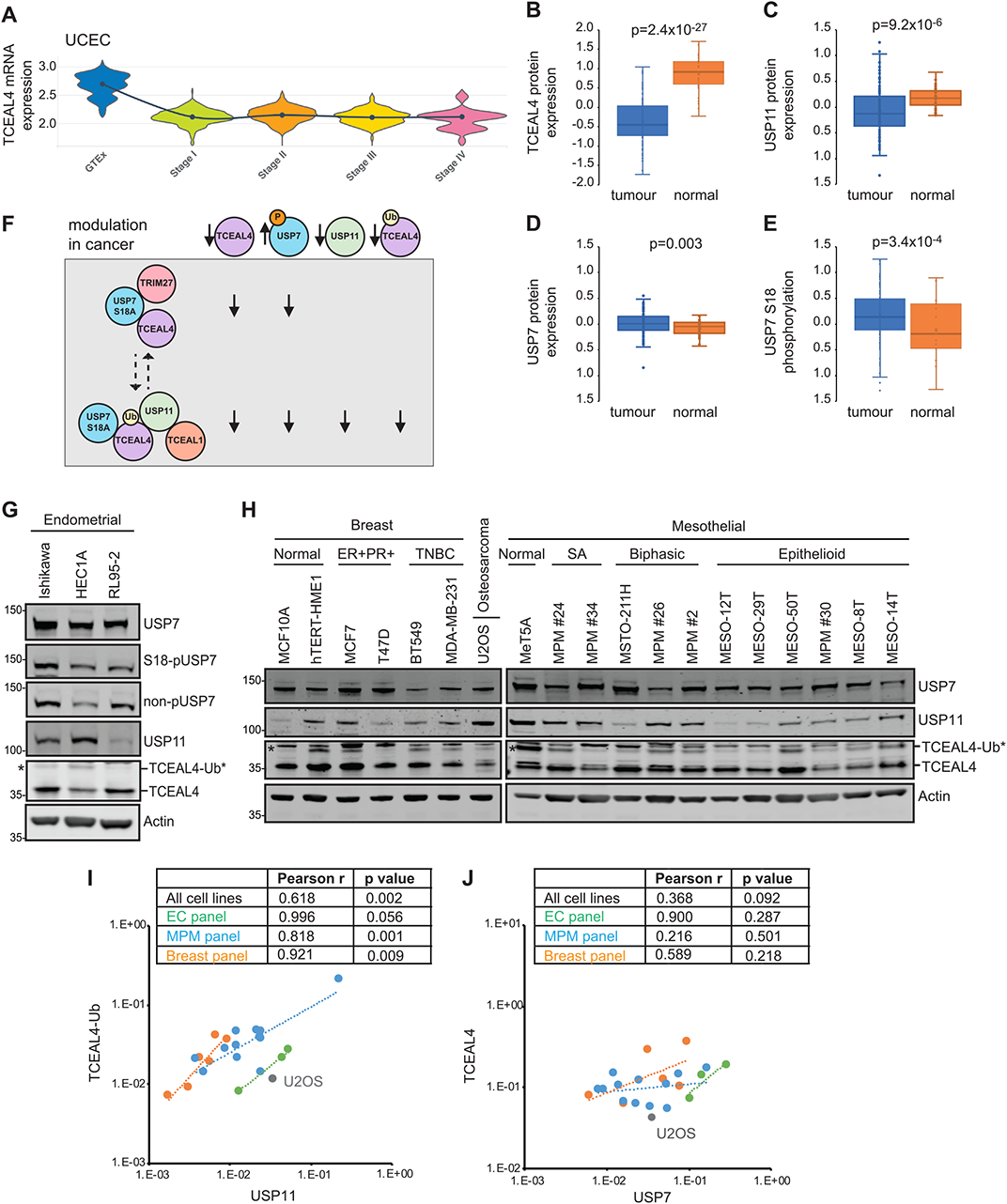
TCEAL4/USP7 complex components are modified in cancer. **A**, TCEAL4 gene expression is downregulated in uterine corpus endometrial cancer (UCEC). Violin plot showing TCEAL4 gene expression (log_2_ Z-score) by AJCC clinical grade with trendline of median values; GTEx, normal tissue. **B-E**, TCEAL4 (**B**) and USP11 (**C**) protein expression are reduced whilst USP7 expression (**D**) and S18-phosphorylation (**E**) are increased in UCEC. CPTAC global proteomics or phosphoproteomics site level data (log_2_ transformed); t-test for tumour (blue) versus age-matched normal tissue (orange). **F**, Model proposing how S18-dephosphorylated USP7 complexes might be altered in cancer. **G & H**, Immunoblots for USP7 complex components in panels of cell lines representing endometrial cancer, breast cancer and mesothelioma. **I & J**, USP11 expression positively correlates with TCEAL4-Ub levels. Quantification of immunoblots shown in (**G**) and (**H**).

To explore these relationships in more detail and examine levels of the two different TCEAL4 species we used a panel of cell lines representing endometrial cancer, as well as breast cancer and mesothelioma (Figure 8G & 8H). TCEAL4-Ub levels varied, generally being lower in normal cells or less aggressive tumours. Notably, TCEAL4-Ub positively correlated with USP11 protein expression across the cell panel (Figure 8I) consistent with our observation that USP11 is required to maintain TCEAL4-Ub levels. Correlation of TCEAL4 with total USP7 expression was less marked (Figure 8J), but this tracked closely with S18-unphosphorylated USP7 in endometrial cancer cells (Figure 8G), consistent with our initial finding that TCEAL4 preferentially interacts with non-phosphorylatable USP7 (Figure 3). Therefore, TCEAL4/S18-dephosporylated USP7 complexes could also be modulated in cancer through downregulation of TCEAL4-Ub, which is dependent upon USP11 expression (Figure 8F).

In summary, we describe a novel role for S18-phosphorylation in biasing the cellular interactions of USP7, such that non-phosphorylated USP7 interacts with TCEAL4 and USP11, which are both USP7 substrates. We also report that USP11 specifically interacts with and protects a mono-ubiquitylated TCEAL4 species, which in turn can scaffold a complex with USP7. Overall, we provide evidence for distinct complexes with the potential to direct cellular USP7 activity that may be modulated in cancer.

## Discussion

To enable diverse roles through alternative substrates, USP7 must be tightly regulated within cells. However, understanding of USP7 recruitment into specific complexes that may direct cellular activity remains incomplete. Here we show that reversible S18-phosphorylation of USP7 is one driver and, for S18-dephosphorylated USP7, that selectivity is further influenced by alternative complexes with the previously uncharacterised protein TCEAL4. Serine-18 resides in the unstructured N-terminal of USP7, adjacent to the TRAF domain where p53, MDM2, USP11 and PPM1G interact via P/A/ExxS motifs ^35, 45^. Although there is no full-length USP7 structure, models based on specific domains and partial structures ^14, 17, 46^ position the N-terminal alongside the catalytic domain, and potentially close to the second ubiquitin-like (UBL) domain, which binds proteins such as ICP0 and GMPS with KxxxKxK/R motifs ^16, 47^. Thus S18-phosphorylation could conceivably modulate USP7 catalytic activity or its interaction with specific substrates or adaptors; we found no evidence for altered DUB activity, but rather that it influences the palette of proteins with which USP7 interacts.

We identified several USP7 interactors in U2OS cells that were reported in HeLa or AGS cells ^8, 35, 48^ and other novel interactors, such as the paralogs MAGED1 and MAGED4. Whilst some proteins like MAGED4 were detected at similar levels irrespective of USP7-S18 mutation, others were enriched by either S18E or S18A mutants, with MAGED1 showing preference for phospho-mimetic USP7-S18E. MAGE proteins are common interaction partners for E3 RING ubiquitin ligases and can enhance their catalytic activity ^49^. Notably, MAGED1 interacts with the RING ligase PJA1 ^49^ that was also preferentially pulled down with S18E-USP7. MAGED1 may activate PJA1 yet is also its substrate ^34^ thus S18-phosphorylated USP7 could potentially stabilise a MAGED1/PJA1 complex as a molecular rheostat, analogous to the role for USP7 in the MAGEL2/TRIM27 complex ^15^.

Here however, we focused on delineating TCEAL/USP11 complexes that preferentially form with phospho-null USP7-S18A, as a model for the stress-responsive form of USP7. In DUB-orientated studies, USP11 and TCEAL4 were previously described as USP7 interactors ^35, 48^ whilst TCEAL1 was validated as a USP11 interactor ^8^; indeed we show that USP11 is required to bring TCEAL1 into a complex with USP7-S18A. Interrogation of a multi-dimensional interactome for HeLa cells confirms a GFP-USP11 complex with TCEAL1, TCEAL4 and USP7 ^50^. In that study, GFP-TCEAL4 interacted with USP7, USP11 and TRIM27, however their stoichiometry predicted that only TCEAL4 and USP11 form a stable core complex, with USP7 excluded because of higher abundance stoichiometry ^50^. This may reflect participation of USP7 in multiple cellular complexes, and our finding that TCEAL4 preferentially interacts with USP7-S18A suggests a limited cellular pool of S18-dephosphorylated USP7 is actually available to participate in core TCEAL4 complexes. Indeed, we saw robust reciprocal interaction of USP7 and TCEAL4 in U2OS cells, even when USP11 was depleted. Although the nature of the interaction between USP7 and TCEAL4 remains to be characterised, TCEAL4 has predicted P/A/ExxS and KxxxKxK motifs (Supplementary Figure S6), which could potentially mediate USP7 binding. Reciprocal regulation of DUBs and their interaction partners are common features of regulatory networks. Here we found that USP11 and USP7 co-ordinately regulate stability of TCEAL1 and TCEAL4, whilst speculating that alternative TCEAL4 complexes might support specific cellular roles of USP7.

We find that TCEAL4 exists as two species that are incorporated into different USP7 complexes. It is unclear why the most abundant lower MW TCEAL4 species migrates abnormally on gel electrophoresis, although we speculated this might be consistent with SUMO-1 modification. However, constitutive SUMOylation is rare, and we were unable to confirm any candidate SUMO-modification sites. However, we demonstrated that the less stable higher MW TCEAL4-Ub species is primarily ubiquitylated at K142. The approximately 8kDa shift for TCEAL4-Ub relative to TCEAL4 is consistent with mono-ubiquitylation, although the approximately 17kDa shift compared to the predicted MW for unmodified TCEAL4 could indicate two ubiquitin moieties. However, if the latter were the case, K142 mutation did not yield a product indicative of multiple mono-ubiquitylation, suggesting any second mono-ubiquitylation site is dependent upon K142 modification. Indeed, three ubiquitylated TCEAL4 lysines (K139, K142 and K164) were recently identified in HCT116 cells, with 5-fold higher basal occupancy at K142, although these likely represent poly-ubiquitylation ^51^.

Our data show that USP11 is necessary but not sufficient for the presence of TCEAL4-Ub, and that USP11 is only found in complex with the TCEAL4-Ub species. Intriguingly, TCEAL4 mono-ubiquitylation is preserved within the complex and not removed by USP11 DUB activity. Whilst it is possible that USP11 may edit ubiquitin chains on TCEAL4 there is no prior evidence that USP11 cannot remove mono-ubiquitin, despite a preference for K63/K6/K33/K11 over other ubiquitin chains ^52^. Alternatively, an E2/E3 ubiquitin ligase may be recruited within the USP11 complex that generates TCEAL4-Ub. Indeed, TCEAL4-Ub is not detected in HCT116 cells despite USP11 expression, potentially reflecting the lack of such a ubiquitin ligase in this cell line.

We describe a complex interplay between USP7 and USP11 in regulating the two TCEAL4 species, with USP7 acting as the primary DUB (Figure 7D). When both USP11 and USP7 are present, TCEAL4-Ub scaffolds a complex in which USP7, like USP11, does not remove the K142-monoubiquitin. However, when USP11 levels are low, or its regulation of TCEAL4-Ub is decoupled as in HCT116 cells, USP7 appears to remove both the mono-ubiquitin and poly-ubiquitin chains from TCEAL4. Recent proteomic studies support our finding that TCEAL4 is a USP7 substrate, as inhibition of USP7 with XL177A in MM.1S myeloma cells, or FT671 in HCT116 cells, reduced TCEAL4 expression ^51, 53^ and increased TCEAL4 poly-ubiquitylation at K139, K142 and K164 ^51^. We find that USP7 also stabilises other components of core TCEAL4 complexes, consistent with previous reports that USP7 protects TRIM27 from auto-ubiquitylation and proteasomal degradation ^15^ and that USP11 may be a context-dependent USP7 substrate ^35^. Here we show that USP11 is ubiquitylated and destabilised upon USP7 depletion in U2OS cells, but not in HCT116 cells, and speculate this context-dependence relies upon TCEAL4-Ub, which when present can scaffold interaction between USP7 and USP11.

The molecular functions and physiological importance of TCEAL4 are unknown, however our interactome suggests a variety of cellular roles for TCEAL4 and provides a valuable resource for further investigation. For example, TCEAL4 interacts with proteins involved in responses to cellular stress, including DNA repair, consistent with the notion of S18-dephosphorylated USP7 as a stress-responsive form of the DUB. Interestingly, the common interactors of TCEAL4, USP7 and USP11 that we identified include the spindle assembly checkpoint proteins MAD2, BUB3, RAE1 and the dynein/dynactin complex, which may point towards roles in mitotic cell division for the TCEAL4-Ub scaffolded complex. Indeed, BUB3 and RAE1 are known interactors of both DUBs ^8^ and validated substrates of USP7 and USP11, respectively ^54, 55^. It will therefore be of interest in future to investigate roles for the specific TCEAL4/USP7 complexes, both in the context of cellular stress and during cell division.

Amongst the TCEAL4 family, TCEAL9 is a candidate oncogene ^56, 57^ whilst TCEAL7 is a putative tumour-suppressor ^39, 40^. TCEAL4 is downregulated in anaplastic thyroid cancer ^58^, suggesting a tumour suppressive role consistent with transcriptomic data for ovarian and endometrial cancer. However, TCEAL4 may also be dysregulated in cancer through inter-conversion of the two cellular species. We show that amongst a panel cancer cell lines, including those with or without evidence for downregulated TCEAL4 transcription, TCEAL4-Ub levels are variable and positively correlate with USP11 expression. This may influence the balance between USP7 complexes with TCEAL4-Ub or TCEAL4, particularly where USP7 stabilises USP11 with the potential to drive a feedforward loop towards the TCEAL4-Ub complex. With continuing development of highly specific USP7 inhibitors, understanding how S18-phosphorylation and TCEAL4 complexes contribute to the context-specificity and dynamics of USP7 function will be important to rationally exploit USP7-directed cancer therapy.

## Methods

### Plasmids, cloning and mutagenesis

USP7 (NM_003470.2) was cloned from pEGFP-USP7 (gift from Prof Sylvie Urbe, Liverpool, UK) into pCR4-TOPO (Thermo Fisher Scientific). Catalytically inactive and phospho-mutant forms of USP7 were generated by Quickchange site-directed mutagenesis in pCR4-TOPO using complementary primer pairs, forward primers: USP7(C223S) 5’-GAATCAGGGAGCGACT<u>TCT</u>TACATGAACAGCCTGC-3’; USP7(S18A) 5’-CGAGCAGCAGTTG<u>GCC</u>GAGCCCGAGGAC-3’; USP7(S18E) 5’-CGAGCAGCAGTTGAGCG<u>GAG</u>CCGAG GACATGGAGATG-3’. pCR4-TOPO USP7 plasmids were sequence verified and shuttled into the destination vector pEGFP_GW (gift from Prof Sylvie Urbe) using the Gateway system (Invitrogen, Thermo Fisher Scientific). GFP or GFP-tagged USP7 WT and mutants were then subcloned into the pcDNA5/FRT/TO vector (Invitrogen, Thermo Fisher Scientific) and sequence verified before generating inducible FlpIn T-Rex cell lines. pCMV6-TCEAL4-MYC-FLAG isoform-1 plasmid (CAT#: RC214906, NM_024863, human TCEAL4 transcript variant 1) and pCMV6-TCEAL4 isoform-2 (CAT#: SC335597, NM_001300901, human TCEAL4 transcript variant 5) were both from Origene. TCEAL4 isoform-1 was cloned from pCMV6-TCEAL4-MYC-FLAG with the Gateway cloning system into pDONR223 and then into the destination vector pEGFP_GW. K142R TCEAL4 mutant was generated by Quickchange site-directed mutagenesis in pEGFP_GW using complementary primer pairs, forward primer: 5’-TACCTCAAGGAGTAT<u>AGA</u>GAGGCTATACATGATATG-3’, and sequence verified. pCMV2-FLAG-Ub was a gift from Dr John O’Bryan to Prof Urbe’s lab.

### Cell culture

A549 (ECACC) and U2OS (ATCC) cells were cultured in Dulbecco’s modified Eagle’s medium (DMEM) supplemented with 10% fetal bovine serum (FBS) and 1% non-essential amino acids (NEAA). MeT5A cells (ATCC) were cultured in supplemented Medium-199 and mesothelioma cell lines (ATCC or MesobanK ^59^) in RPMI-1640 supplemented with 10% FBS as described elsewhere (Barnett *et al* & Butt *et al*, manuscripts in preparation). MCF10A (Horizon Discovery) were cultured as previous described ^60^. hTERT-HME1 (ATCC) and breast cancer cell lines (ATCC or local collaborators) were cultured in recommended media as described elsewhere (Sabat-Pospiech *et al*, manuscript in preparation). Endometrial cancer lines were sourced and cultured as described previously ^61^. All cells were cultured in a humidified incubator at 37°C and 5% CO_2_. Cell lines were authenticated by STR profiling, were mycoplasma-free and were cultured for limited passages.

### Inducible cell lines

For generation of stable U2OS lines, pcDNA5/FRT/TO USP7 plasmids were co-transfected with pOG44 (Invitrogen, Thermo Fisher Scientific) into inducible FlpIn T-Rex U2OS cells (gift from Prof Steven C. Blacklow, Boston, MA, USA), using GeneJuice (Merck Millipore). Stable cell lines were selected with 150 µg/ml hygromycin and 15 µg/ml blasticidin (Thermo Fisher Scientific). Individual cell clones were induced with 1µg/ml doxycycline for 24 hours and expression of the USP7 proteins assessed by immunoblotting and fluorescence microscopy. For the selected clones, genomic DNA was extracted using the Blood & Cell Culture DNA Kit (Qiagen) and the GFP-USP7 insert was PCR amplified for sequence verification. U2OS FlpIn T-Rex cell lines were cultured in DMEM supplemented with 10% tetracycline free-FBS (LabTech) and 1% NEAA; they were treated with 1µg/ml doxycycline for 24 hours prior to analysis to induce the expression of GFP or GFP-USP7 proteins.

### siRNA and plasmid transfection

U2OS cells were seeded at appropriate density and transfected the following day with 10nM (unless otherwise specified) siRNA using Lipofectamine RNAiMax (Life Technologies) for 72 hrs, and/or with plasmid DNA (e.g. 1µg/well for 6-well plates) for 24 hrs using GeneJuice (Merck Millipore). siRNAs targeting USP7, USP11, TCEAL4 and TCEAL2 were siUSP7_1 (J-006097-05, Dharmacon), siUSP7_3 (J-006097-07, Dharmacon), siUSP11_5 (J-006063-05, Dharmacon), siUSP11_8 (J-006063-08, Dharmacon), siTCEAL4_6 (SI04311293, Qiagen), siTCEAL4_7 (SI04312721, Qiagen), siTCEAL4_23 (J-016799-23, Dharmacon), siTCEAL4_25 (J-016799-25, Dharmacon), siTCEAL2_19 (J-015344-19, Dharmacon) and siTCEAL2_20 (J-015344-20, Dharmacon). Non-targeting siRNA controls were siC (Qiagen All-Stars negative control,1027281) or siNT1 (D-001810-10-05, Dharmacon).

### Antibodies

Antibodies used in this study were: mouse anti-β-actin (ab6276, Abcam), rabbit anti-actin (A2066, Sigma-Aldrich), anti-USP7 (ab4080, Abcam), anti-S18 phosphorylated USP7 (ABC225, Merck-Millipore), anti-S18 non-phosphorylated USP7 (ABC226, Merck-Millipore), anti-USP11 (A301-613A, Bethyl Laboratories), anti-TRIM27 (12205-1-AP, Proteintech, or NBP2-46214, Novus Biologicals), anti-TCEAL4 (SAB1400709, Sigma-Aldrich, or PA567536, Invitrogen, Thermo Fisher Scientific), anti-TCEAL1 (sc-393621, Santacruz), anti-p53 (sc-126, Santacruz), anti-FLAG (F3165 or F7425, Sigma-Aldrich), anti-Ub (PW8810, Enzo Life Sciences) and anti-SUMO1 (#4930, Cell Signaling, or 33-2400, Invitrogen). Polyclonal affinity-purified sheep anti-GFP was a gift from Ian Prior (Liverpool, UK).

### Inhibitors

To accumulate ubiquitylated proteins, cells were incubated in 50nM epoxomicin (Merck-Millipore) for 6 hrs where indicated. The USP7 inhibitor XL177A ^29^ and its 500-fold less potent enantiomer XL177B were a gift from Dr Sara Buhrlage (Dana-Farber Cancer Institute, Boston). U2OS cells were incubated with 10μM XL177A or XL177B for 6 hrs.

### Immunoblotting

Whole cell extracts were prepared by direct addition of hot Laemmli buffer and incubation at 110°C for 10min with intermittent vortexing. Protein concentrations were determined by Bicinchoninic Acid (BCA) assay (Thermo Scientific). Following resolution by SDS-PAGE, proteins were transferred to BiotraceNT membrane (VWR) and incubated with primary antibodies. Proteins were visualized using donkey anti-mouse, anti-rabbit or anti-sheep secondary antibodies conjugated to the IRDyes IR680-LT, or IR800 (LI-COR) and a LI-COR Odyssey system, with images quantified in ImageStudio.

### Immunoprecipitation

Cells were lysed in NP-40 lysis buffer (0.5% NP-40, 25mM Tris pH 7.5, 100mM NaCl, 50mM NaF, 2mM MgCl_2_, 1mM EGTA, containing protease inhibitors and PhosStop (Sigma-Aldrich) for 20 minutes on ice, then pre-cleared by centrifugation. Lysates were adjusted to 1 mg/ml in lysis buffer and subject to overnight immunoprecipitation at 4°C with either USP7 or USP11 antibody and protein A agarose beads (Sigma-Aldrich), or with GFP-NanoTrap beads (prepared in house) to pull-down GFP-tagged proteins. Immunoprecipitates were washed three times with wash buffer 1 (0.5% NP-40, 25mM Tris pH 7.5, 100mM NaCl, 50mM NaF, 2mM MgCl_2_) and once in wash buffer 2 (10 mM Tris pH 7.5, 2mM MgCl_2_) before elution in 3X sample buffer (5 mins at 95°C) and analysis by immunoblotting. For mass spectrometry protein elution from GFP-NanoTrap beads was performed twice with 50μl of elution buffer (1% SDS, 1% β-mercaptoethanol, 10mM Tris pH 6.8).

### Mass Spectrometry

For SILAC experiments, cells were grown and maintained in arginine and lysine free DMEM and 10% tetracycline free-FBS (Labtech) supplemented with the following amino acids: “Light” cells; l-lysine (Lys0), l-arginine (Arg0) and l-proline (Pro0). “Medium” cells; l-lysine-2H4 (Lys4), l-arginine-U–13C6 (Arg6) and l-proline (Pro0). “Heavy” cells; l-lysine-U–13C6-15N2 (Lys8), l-arginine-U–13C6–15N4 (Arg10) and l-proline (Pro0). Final concentrations of amino acids were 84 mg/L arginine, 146 mg/L lysine and 200mg/L Proline. Immunoprecipitated proteins were eluted and 50µg (5%) of each IP, 10µg input and 10µg flow-through (FT) were analysed by immunoblotting with a GFP antibody. The elutes were reduced (5 mM DTT, 60 °C, 25 min) and alkylated (20 mM iodoacetamide, 30 min, room temperature) and samples concentrated with Amicon Ultra-4 columns (Merck-Millipore), After protein precipitation with methanol/chloroform, samples were incubated with mass spectrometry grade Trypsin Gold (Promega) for 16h at 37°C. Peptides were extracted with ethyl acetate and desalted on C18-Stage tip columns (Thermo Fisher Scientific), eluates were dried in a SpeedVac and resuspended in 1% formic acid for loading. LC-MS/MS was performed by the Proteomics Research Technology Platform (University of Warwick) with reverse phase chromatography to separate peptides using an Ultimate 3000 RSLCnano system (Dionex) before analysis on a Thermo Orbitrap Fusion (Q-OT-qIT, Thermo Scientific). RAW files from LC-MS/MS were processed using MaxQuant version 1.5.3.8 or 1.6.7.0 (http://www.maxquant.org) and the UniProt human protein database for protein identification.

### Ubiquitin active site-directed probe assays

Cells were lysed in non-denaturing buffer (50mM Tris pH7.5, 5mM MgCl2, 250mM sucrose, 1mM DTT, 2mM ATP) on ice and homogenized by progressively passing through 23G, 26G and 30G needles, PhosStop (Sigma-Aldrich) was added immediately afterwards. Lysates were cleared by centrifugation (20 min, 14,000 rpm, 4°C) and a BCA assay performed to standardize protein concentrations. 10μg protein from U2OS USP7 FlpIn T-Rex cells, induced with doxycycline, was incubated with 50ng HA-Ub-VME (UbiQ-035, UbiQ, Amsterdam, The Netherlands) at a 1:200 ratio of probe:protein, for 5 or 15 min at 37°C with shaking at 300rpm. 15μg protein from A549 cells was incubated with 37.5 ng HA-Ub-VME (1:400 ratio) for 2, 5, 10, 15 or 15 min. 5x sample buffer (15% SDS, 312.5mM Tris pH6.8, 50% glycerol, 16% β-mercaptoethanol, bromophenol blue) was used to terminate the reaction at 95°C for 5 min before SDS-PAGE analysis.

### USP7 protein expression and purification

A codon-optimised sequence for full-length USP7 (USP7FL) was cloned into pGEX6p-1 using BamHI/NotI restriction sites (Addgene, #63573). Mutations at S18 were introduced using partially overlapping primers with Phusion Flash polymerase (Thermo Fisher Scientific); all clones were sequence-verified. USP7FL constructs were grown in *Escherichia coli* BL21 Rosetta2 (DE3) in 0.5 L Terrific Broth at 37°C to OD_600_ of 1.8-2.0, then rapidly cooled and induced with 100 μM IPTG overnight at 18°C. Cultures were harvested and resuspended in 50 mM HEPES pH 7.5, 250 mM NaCl, 1 mM EDTA, 1 mM DTT with USP7 proteins purified as described previously ^62^ using Glutathione Sepharose 4B beads (GE Healthcare). After elution the GST tag was removed under dialysis using 3C protease and the sample fractionated by PorosXQ anion exchange (Thermo Fisher Scientific). Appropriate fractions were pooled, concentrated and further purified on a Superdex gel filtration column (GE Healthcare). The peak fractions were pooled, concentrated to 1 mg/mL and flash frozen.

### USP7 *in vitro* enzyme activity assay

Wildtype and mutant USP7FL were assessed for activity using the minimal substrate Ubiquitin-Rhodamine (UbRho; UbiQ, the Netherlands), by measuring the fluorescence intensity upon cleavage of the rhodamine group. The enzyme was diluted to 2 nM in running buffer (20 mM HEPES 7.5, 100 mM NaCl, 1 mM DTT and 0.05% Tween-20); a titration range of UbRho was prepared in the same buffer. The two components were combined 1:1 in a final volume of 20 µL in and the reaction monitored in a Pherastar plate reader (BMG LABTECH GmbH, Germany) using excitation at 485 nm (±10 nm) and detection at 520 nm (±10 nm). The fluorescence intensity readings were converted into product concentrations and the initial velocity was determined through linear regression of the initial slope using Prism 7 (GraphPad). The slopes were then plotted against the substrate concentration and Prism 7 regression for Michaelis-Menten used to determine the V_max_ and K_M_.

### Bioinformatics

TCEAL4 gene expression data for uterine corpus endometrial cancer (UCEC) was accessed via canSAR (https://cansarblack.icr.ac.uk). CPTAC global proteomics or phosphoproteomics site level data (median polishing, log_2_ transformed) of interest was extracted from supplementary data for reference ^44^. TCEAL4 interacting proteins were analysed using the gene set enrichment analysis (GSEA) software with the molecular signatures database (MSigDB) collection of annotated gene-sets. MSigDB C2 collection (curated gene sets), including curated pathways (CP), REACTOME, Biocarta and KEGG pathways, was used for the enrichment analysis. TCEAL4 interactors potentially involved in DNA repair were assigned by a combination of manual annotation from the literature and bioinformatics using GSEA/MSigDB.

### Statistics

All statistical tests for experimental data were performed using GraphPad Prism version 6.00/8.4.1 for Mac; P values less than 0.05 were considered to be significant. Data were analysed by T-test or ANOVA, as appropriate. Statistical tests and n numbers are indicated in figure legends. These parametric tests are suitable for continuous data sets without off-scale measurements, and assume Gaussian distribution, which was confirmed by the Shapiro-Wilk normality test in Prism, and equivalent variance between samples confirmed by a variance homogeneity test.

## Data availability

All data generated or analysed during this study are included in this published article (and its supplementary information files).

## Acknowledgements

This work was funded by North West Cancer Research, project grant reference CR1093 (JMC, FQ). SD and ICW were supported on studentships funded by Wellcome, grant references 096567/Z/11/Z SD) and 205806/Z/16/Z (ICW). RQK and TKS were funded by the Dutch Cancer Society project 2012-5398 and the Oncode Institute. We thank Dr Chris Hill (DKH lab) for providing lysates from endometrial cancer cell lines.

This research was funded in whole, or in part, by the Wellcome Trust [*096567/Z/11/Z and 205806/Z/16/Z*]. **For the purpose of Open Access, the author has applied a CC BY public copyright licence to any Author Accepted Manuscript version arising from this submission**.

## Author Contributions

F.Q. designed and performed experiments unless otherwise indicated, analysed data, performed bioinformatic analyses, and contributed to writing the manuscript. S.D. designed and generated the USP7 Flp-In cell lines and contributed to initial experiments on USP7 cellular activity and localisation. R.Q.K. designed and performed *in vitro* USP7 activity assays. I.C.W. designed and performed some TCEAL4 mutagenesis, TCEAL4-GFP imaging and TCEAL4 isoform RT-PCR. D.K.H. advised on endometrial cancer and cell line analyses. T.K.S. contributed to *in vitro* experiments, data interpretation and provided critical feedback. J.M.C. conceived and supervised the study, interpreted data and wrote the manuscript. All authors discussed the results and contributed to the manuscript.

## Materials & Correspondence

Correspondence and material requests should be addressed to J.M.C.

## References

1. Grabbe C, Husnjak K, Dikic I. The spatial and temporal organization of ubiquitin networks. Nat Rev Mol Cell Biol 12, 295–307 (2011).

2. Clague MJ, Coulson JM, Urbé S. Cellular functions of the DUBs. J Cell Sci 125, 277–286 (2012).

3. Clague MJ, Urbe S, Komander D. Breaking the chains: deubiquitylating enzyme specificity begets function. Nat Rev Mol Cell Biol 20, 338–352 (2019).

4. Heideker J, Wertz IE. DUBs, the regulation of cell identity and disease. Biochem J 467, 191 (2015).

5. Sacco JJ, Coulson JM, Clague MJ, Urbe S. Emerging roles of deubiquitinases in cancer-associated pathways. IUBMB Life 62, 140–157 (2010).

6. Sahtoe DD, Sixma TK. Layers of DUB regulation. Trends Biochem Sci 40, 456–467 (2015).

7. Rawat R, Starczynowski DT, Ntziachristos P. Nuclear deubiquitination in the spotlight: the multifaceted nature of USP7 biology in disease. Curr Opin Cell Biol 58, 85–94 (2019).

8. Sowa ME, Bennett EJ, Gygi SP, Harper JW. Defining the human deubiquitinating enzyme interaction landscape. Cell 138, 389–403 (2009).

9. Li M, et al. Deubiquitination of p53 by HAUSP is an important pathway for p53 stabilization. Nature 416, 648–653 (2002).

10. Song MS, et al. The deubiquitinylation and localization of PTEN are regulated by a HAUSP-PML network. Nature 455, 813–817 (2008).

11. Li M, Brooks CL, Kon N, Gu W. A dynamic role of HAUSP in the p53-Mdm2 pathway. Mol Cell 13, 879–886 (2004).

12. Jin Q, et al. USP7 Cooperates with NOTCH1 to Drive the Oncogenic Transcriptional Program in T-Cell Leukemia. Clin Cancer Res 25, 222–239 (2019).

13. Darling S, Fielding AB, Sabat-Pospiech D, Prior IA, Coulson JM. Regulation of the cell cycle and centrosome biology by deubiquitylases. Biochem Soc Trans 45, 1125–1136 (2017).

14. Kim RQ, Sixma TK. Regulation of USP7: A High Incidence of E3 Complexes. J Mol Biol 429, 3395–3408 (2017).

15. Hao YH, et al. USP7 Acts as a Molecular Rheostat to Promote WASH-Dependent Endosomal Protein Recycling and Is Mutated in a Human Neurodevelopmental Disorder. Mol Cell 59, 956–969 (2015).

16. Faesen AC, Dirac AM, Shanmugham A, Ovaa H, Perrakis A, Sixma TK. Mechanism of USP7/HAUSP Activation by Its C-Terminal Ubiquitin-like Domain and Allosteric Regulation by GMP-Synthetase. Mol Cell 44, 147–159 (2011).

17. Rouge L, et al. Molecular Understanding of USP7 Substrate Recognition and C-Terminal Activation. Structure 24, 1335–1345 (2016).

18. Kim RQ, et al. Kinetic analysis of multistep USP7 mechanism shows critical role for target protein in activity. Nat Commun 10, 231 (2019).

19. van der Knaap JA, et al. GMP synthetase stimulates histone H2B deubiquitylation by the epigenetic silencer USP7. Mol Cell 17, 695–707 (2005).

20. Morotti A, et al. BCR-ABL disrupts PTEN nuclear-cytoplasmic shuttling through phosphorylation-dependent activation of HAUSP. Leukemia 28, 1326–1333 (2014).

21. Khoronenkova SV, Dianova, II, Ternette N, Kessler BM, Parsons JL, Dianov GL. ATM-dependent downregulation of USP7/HAUSP by PPM1G activates p53 response to DNA damage. Mol Cell 45, 801–813 (2012).

22. Tang J, Qu L, Pang M, Yang X. Daxx is reciprocally regulated by Mdm2 and Hausp. Biochem Biophys Res Commun 393, 542–545 (2010).

23. Tang J, et al. Critical role for Daxx in regulating Mdm2. Nat Cell Biol 8, 855–862 (2006).

24. Reddy BA, et al. Nucleotide biosynthetic enzyme GMP synthase is a TRIM21-controlled relay of p53 stabilization. Mol Cell 53, 458–470 (2014).

25. Harrigan JA, Jacq X, Martin NM, Jackson SP. Deubiquitylating enzymes and drug discovery: emerging opportunities. Nat Rev Drug Discov, (2017).

26. Turnbull AP, et al. Molecular basis of USP7 inhibition by selective small-molecule inhibitors. Nature 550, 481–486 (2017).

27. Gavory G, et al. Discovery and characterization of highly potent and selective allosteric USP7 inhibitors. Nat Chem Biol 14, 118–125 (2018).

28. Ohol YM, et al. Novel, Selective Inhibitors of USP7 Uncover Multiple Mechanisms of Antitumor Activity In Vitro and In Vivo. Mol Cancer Ther 19, 1970–1980 (2020).

29. Schauer NJ, et al. Selective USP7 inhibition elicits cancer cell killing through a p53-dependent mechanism. Sci Rep 10, 5324 (2020).

30. Kon N, Kobayashi Y, Li M, Brooks CL, Ludwig T, Gu W. Inactivation of HAUSP in vivo modulates p53 function. Oncogene 29, 1270–1279 (2010).

31. Kon N, et al. Roles of HAUSP-mediated p53 regulation in central nervous system development. Cell Death Differ, (2011).

32. Faesen AC, et al. The differential modulation of USP activity by internal regulatory domains, interactors and eight ubiquitin chain types. Chem Biol 18, 1550–1561 (2011).

33. Fernandez-Montalvan A, et al. Biochemical characterization of USP7 reveals post-translational modification sites and structural requirements for substrate processing and subcellular localization. FEBS J 274, 4256–4270 (2007).

34. Teuber J, Mueller B, Fukabori R, Lang D, Albrecht A, Stork O. The ubiquitin ligase Praja1 reduces NRAGE expression and inhibits neuronal differentiation of PC12 cells. PLoS One 8, e63067 (2013).

35. Georges A, Marcon E, Greenblatt J, Frappier L. Identification and Characterization of USP7 Targets in Cancer Cells. Sci Rep 8, 15833 (2018).

36. Winter EE, Ponting CP. Mammalian BEX, WEX and GASP genes: coding and non-coding chimaerism sustained by gene conversion events. BMC Evol Biol 5, 54 (2005).

37. Navas-Perez E, et al. Characterization of an eutherian gene cluster generated after transposon domestication identifies Bex3 as relevant for advanced neurological functions. Genome Biol 21, 267 (2020).

38. Hornbeck PV, Zhang B, Murray B, Kornhauser JM, Latham V, Skrzypek E. PhosphoSitePlus, 2014: mutations, PTMs and recalibrations. Nucleic Acids Res 43, D512–520 (2015).

39. Chien J, et al. A role for candidate tumor-suppressor gene TCEAL7 in the regulation of c-Myc activity, cyclin D1 levels and cellular transformation. Oncogene 27, 7223–7234 (2008).

40. Rattan R, et al. TCEAL7, a putative tumor suppressor gene, negatively regulates NF-kappaB pathway. Oncogene 29, 1362–1373 (2010).

41. Hendriks IA, Lyon D, Young C, Jensen LJ, Vertegaal AC, Nielsen ML. Site-specific mapping of the human SUMO proteome reveals co-modification with phosphorylation. Nat Struct Mol Biol 24, 325–336 (2017).

42. Lumpkin RJ, et al. Site-specific identification and quantitation of endogenous SUMO modifications under native conditions. Nat Commun 8, 1171 (2017).

43. Hendriks IA, et al. Site-specific characterization of endogenous SUMOylation across species and organs. Nat Commun 9, 2456 (2018).

44. Dou Y, et al. Proteogenomic Characterization of Endometrial Carcinoma. Cell 180, 729–748 e726 (2020).

45. Sheng Y, et al. Molecular recognition of p53 and MDM2 by USP7/HAUSP. Nat Struct Mol Biol 13, 285–291 (2006).

46. Hu M, Gu L, Li M, Jeffrey PD, Gu W, Shi Y. Structural basis of competitive recognition of p53 and MDM2 by HAUSP/USP7: implications for the regulation of the p53-MDM2 pathway. PLoS Biol 4, e27 (2006).

47. Pfoh R, et al. Crystal Structure of USP7 Ubiquitin-like Domains with an ICP0 Peptide Reveals a Novel Mechanism Used by Viral and Cellular Proteins to Target USP7. PLoS Pathog 11, e1004950 (2015).

48. Georges A, Coyaud E, Marcon E, Greenblatt J, Raught B, Frappier L. USP7 Regulates Cytokinesis through FBXO38 and KIF20B. Sci Rep 9, 2724 (2019).

49. Doyle JM, Gao J, Wang J, Yang M, Potts PR. MAGE-RING protein complexes comprise a family of E3 ubiquitin ligases. Mol Cell 39, 963–974 (2010).

50. Hein MY, et al. A human interactome in three quantitative dimensions organized by stoichiometries and abundances. Cell 163, 712–723 (2015).

51. Steger M, Ihmor P, Backman M, Müller S, Daub H. Deep ubiquitination site profiling by single-shot data-independent acquisition mass spectrometry. bioRxiv preprint doi: https://doiorg/101101/20200723218651;, (2020).

52. Harper S, Besong TM, Emsley J, Scott DJ, Dreveny I. Structure of the USP15 N-Terminal Domains: A beta-Hairpin Mediates Close Association between the DUSP and UBL Domains. Biochemistry 50, 7995–8004 (2011).

53. Bushman JW, et al. Proteomics-Based Identification of DUB Substrates Using Selective Inhibitors. Cell Chem Biol 28, 78–87 e73 (2021).

54. Stockum A, Snijders AP, Maertens GN. USP11 deubiquitinates RAE1 and plays a key role in bipolar spindle formation. PLoS One 13, e0190513 (2018).

55. Giovinazzi S, et al. Usp7 protects genomic stability by regulating Bub3. Oncotarget 5, 3728–3742 (2014).

56. Gylfe AE, et al. Identification of candidate oncogenes in human colorectal cancers with microsatellite instability. Gastroenterology 145, 540–543 e522 (2013).

57. Ward C, Cauchy P, Garcia P, Frampton J, Esteban MA, Volpe G. High WBP5 expression correlates with elevation of HOX genes levels and is associated with inferior survival in patients with acute myeloid leukaemia. Sci Rep 10, 3505 (2020).

58. Akaishi J, et al. Down-regulation of transcription elogation factor A (SII) like 4 (TCEAL4) in anaplastic thyroid cancer. BMC cancer 6, 260 (2006).

59. Rintoul RC, Rassl DM, Gittins J, Marciniak SJ, Mesoban Kc. MesobanK UK: an international mesothelioma bioresource. Thorax 71, 380–382 (2016).

60. Tang WY, Beckett AJ, Prior IA, Coulson JM, Urbe S, Clague MJ. Plasticity of mammary cell boundaries governed by EGF and actin remodeling. Cell reports 8, 1722–1730 (2014).

61. Kamal A, et al. High AGR2 protein is a feature of low grade endometrial cancer cells. Oncotarget 9, 31459–31472 (2018).

62. Kim RQ, van Dijk WJ, Sixma TK. Structure of USP7 catalytic domain and three Ubl-domains reveals a connector alpha-helix with regulatory role. J Struct Biol 195, 11–18 (2016).

